# A distal CTCF-binding site drives MYC expression plasticity in a negative feed-forward loop

**DOI:** 10.64898/2026.01.28.702300

**Authors:** Chenying Gao, Mirco Martino, Ayushi Chaurasiya, Felipe Beccaria Casagrande, Jia Pei Lim, Yuri Stephan van Ekelenburg, Wei Feng, Yan Liu, Barbara Scholz, Pragyan Mishra, Mallikarjuna Thippana, Dennis Torkornoo, Phong KC Chau, Rabia Khan, Tone Möller Tannaes, Ilias Tzelepis, Ilyas Chacoua, Natalie Geyer, Sebastian Meltzer, Anne H. Ree, Marco Gerling, Johan Hartman, Narsis Kiani, Rolf Ohlsson, Anita Göndör

## Abstract

Oncogene expression heterogeneity systemically diversifies cancer cell phenotypes, enabling selection and cancer evolution. Here we document that the level of MYC expression variation is coordinated by a negative feed-forward loop involving the non-coding *CCAT1* eRNA and the *MYC* gating process. While *CCAT1* eRNA indirectly antagonizes *MYC* gating by promoting *MYC* transcriptional initiation and elongation, the MYC protein promotes gated MYC expression in a feedback loop by inhibiting both *MYC* transcription and *CCAT1* expression. This entire process is coordinated by a CTCF binding site positioned within *CCAT1* at a distal, oncogenic super-enhancer, which functions as a master switch by coordinating both *CCAT1* and gated *MYC* expression to diversify cells toward both low and high MYC levels, as determined by heterogeneity metrics. As hallmarks of this principle are frequent in breast cancer and colorectal tumors, the dynamics of this new principle may underlie transitions between therapy-resistant low-MYC, and proliferative high-MYC tumor cells.

**Highlights:**

- Transcriptional rate regulates *MYC* gating frequency
- *CCAT1* expression indirectly inhibits *MYC* gating by promoting *MYC* transcription
- High MYC protein levels promote gated expression by inhibiting both *MYC* transcription and *CCAT1* expression
- These features are controlled by a single CTCF binding in the distant super-enhancer to drive expression plasticity in colorectal cancer cells

## Introduction

With few exceptions, the pathological activity of the *MYC* gene is a common denominator of cancers, affecting cancer evolution at multiple steps^1^. The encoded protein can function as a repressor or booster of the transcriptional elongation rate of a large subset of target genes to promote the speed of the cell cycle traverse^2-4^, which influences a range of biological processes^5^. The heterogeneity of MYC levels thus pleiotropically influences cell-to-cell variations in phenotypic traits—a feature that can be exploited by tumors to increase their adaptability to changing microenvironments^6^. Previously observed principles of MYC heterogeneity include extrachromosomal ec*MYC* amplification, which is frequent in human cancers^7,8^. As some ec*MYC* copies are distributed stochastically between daughter cells, genotypic heterogeneities may be constantly replenished, thereby providing a feature that might endow cancer cell populations with resistance to hostile environments experienced during metastasis and therapeutic interventions^7,8^. As heterogeneity in MYC protein levels increases tumor plasticity^6^, a well-known feature driving cancer evolution^9,10^, deciphering a detailed mechanistic understanding of how this feature is established, manifested and sustained is essential.

In noncancerous cells, MYC levels are regulated by several layers of negative feedback to prevent pathological MYC expression and thus prevent unscheduled cell proliferation. These include posttranslational modifications regulating the turnover of the MYC protein^11^, the rapid decay kinetics of *MYC* transcripts^12^ and the inhibition of the transcription of its own template with MYC protein overshoot^13^. Pathological expression of the MYC protein can thus be achieved by interfering with any of these processes, including dysregulated signaling pathways and genetic perturbations, such as gene amplification and translocation^2^. Whereas most pathways generating an overactive *MYC* gene require the *MYC* mRNA to be exported out of the nucleus, little is known about the principles regulating this process and if/how it relates to MYC protein heterogeneity.

The *MYC* gene environment is dominated by noncoding genes, such as *CCAT1* that resides within an oncogenic super-enhancer (OSE) positioned approximately 520 kb distal to *MYC*^*14*^. *CCAT1* has received considerable attention because of its ability to promote both *MYC* expression^15,16^ and prevent apoptosis^17^. We have previously shown that the OSE drives the redistribution of active *MYC* alleles to nuclear pores in a WNT signaling-dependent manner in colon cancer cells^18,19^. Proximity to nuclear pores requires interaction between the CTCF binding site within *CCAT1* and the nuclear pore anchor AHCTF1 to dramatically increase the nuclear export rate of the mature *MYC* mRNA, a process termed *MYC* gating^18,19^. Since the decay rate of *MYC* mRNA is several-fold greater in the nucleus than in the cytoplasm, cytoplasmic *MYC* mRNA accumulation is several-fold greater in HCT116 cells than in primary cultures of normal colon epithelial cells lacking *MYC* gating, despite lower levels of *MYC* transcription in HCT116 cells^18^.

We report here that a reduction in *MYC* transcription - triggered by the inhibition or knockdown of CDK9, loss of *CCAT1* expression, or by pathologically increased MYC protein levels - promotes the migration of active but paused *MYC* alleles to nuclear pores, under the control of the colorectal super-enhancer. Following completion of transcription in the nuclear pore microenvironment, MYC protein levels can be rapidly increased in response to extrinsic cues. This dynamic process, termed *MYC* gating, enables cancer cells to simultaneously rescue MYC levels during transcriptional downregulation of *MYC* and increase cell-to-cell heterogeneity of MYC expression, as determined by coefficient of variation, Gini index and cell population redistribution. Thus, the CTCF binding site within the colorectal super-enhancer unleashes dynamic heterogeneity in MYC expression by simultaneously promoting both *CCAT1* expression - that indirectly antagonizes *MYC* gating - and *MYC* recruitment to nuclear pores. Accordingly, a reduction of MYC protein expression to increase its transcriptional rate inhibited the gating process, documenting a feed-forward principle underlying both exponential cell proliferation and expression heterogeneity. Hallmarks of this principle could be documented in both primary and metastatic colorectal (CRC) tumors and breast cancer.

## Results

### CCAT1 eRNA expression antagonizes the cell confluence-dependent gating of MYC to nuclear pores

To address the relationship between *CCAT1* function - a key regulator of *MYC* transcription^16,20^ - and the gating of active *MYC* alleles to nuclear pores, the expression of *CCAT1* eRNA was knocked down by siRNA in HCT116 cells. Since these cells express only the long, nuclear version of the noncoding *CCAT1* eRNA^19^, the cells were subjected to siCCAT1 treatment for three days to achieve a reduction of approximately 70% of its expression (**Fig. 1A)**. In agreement with earlier reports^16,20^, reduction in *CCAT1* expression was accompanied by a reduction of *MYC* mRNA accumulation (**Fig. 1A)** and transcription **(Fig. 1B**).

**Figure 1.**
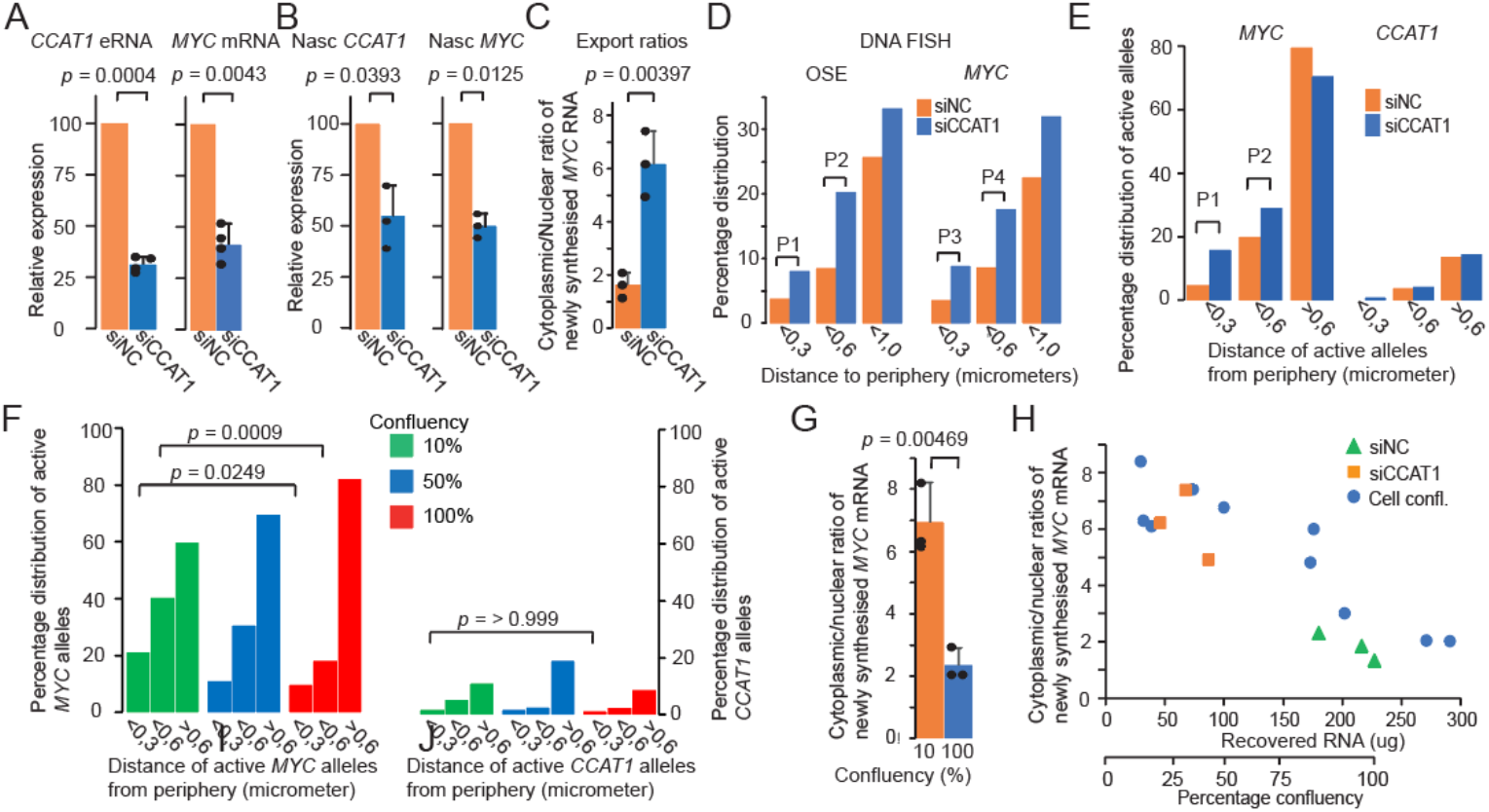
MYC gating is regulated by CCAT1 and cell confluence. (A) si*CCAT1*-mediated reduction in *CCAT1* and *MYC* mRNA accumulation in total RNA samples, normalized first to TBP mRNA accumulation and then to the siNC control samples. (B) siCCAT1-mediated reduction in *CCAT1* and *MYC* transcription in newly synthesized RNA purified from cells labeled with ethynyl-uridine for 15 minutes. The data were first normalized to TBP transcription and then to the siNC control samples. (C) Nuclear export rate analyses comparing the levels of newly synthesized *MYC* transcripts in the cytoplasmic and nuclear fractions. D) Quantitation of DNA FISH signals in relation to the distance to the nuclear periphery. The values were binned from a total of 342 (OSE, siNC), 311 (MYC, siNC), 337 (OSE, siCCAT1) and 306 (MYC, siCCAT1) alleles. P1 = 0.0154; P2 = <0.0001; P3 = 0.0071; P4 = <0.0001. E) Spatial distribution of active *MYC* and *CCAT1* alleles (Stellaris *CCAT1, MYC* exon/intron probes), as determined by RNA FISH, based on the counting of 115 (siNC) and 120 (siCCAT1) alleles. P1 = 0.0087; P2 = 0.124. F) Spatial distribution of active *MYC* and *CCAT1* alleles (Stellaris *CCAT1*/*MYC* exon/intron probes) in cells harvested at 10% (177 alleles),50% (140 alleles) and 100% (132 alleles) confluence, as determined by IncuCyte analyses. The values indicate the binned distance of active *MYC* alleles from the nuclear periphery. G) Nuclear export rates were determined in relation to low and high cell confluence, representing the average of three independent experiments. H) The nuclear export rates were plotted against cell confluence, as determined by the RNA recovery (**Fig. S1C**). The data from panel C is superimposed on this graph to illustrate the relationship between high nuclear export rates and cell confluence, which is affected by the s*i*CCAT1 treatment. All the data are based on three or four (indicated) independent experiments. The *P* values were determined by one-sample (A, B), two-tailed unpaired Student’s t test (C,G) or Fisher’s exact test (D-F). Bar = 10 μm.

We next addressed whether *CCAT1* activity affects the nuclear export rate of processed *MYC* mRNAs by employing our earlier reported strategy to pulse-label HCT116 cells with ethynyl-uridine (EU) for 30 minutes, followed by extensive washes and a 1-hour chase^18^. Following the separation of the nuclear and cytoplasmic fractions, newly synthesized RNAs were isolated and analyzed for the presence of intron 2 and exon 2 *MYC* RNA sequences. Two types of spike-in RNAs enabled the quantitative recoveries of both total nuclear and cytoplasmic RNAs as well as that of EU-labeled newly synthesized RNAs. Contaminations of cytoplasmic and nuclear fractions were quantified by measuring *MYC* intron sequences in the cytoplasm and sequences from the mitochondrial *CYTB* gene in the nucleus^18^. Following normalization *via* spike-in controls and contamination analyses, the results are presented as ratios of cytoplasmic and nuclear exon 2-containing *MYC* RNA. **Figure 1C** shows that treating the cells with siCCAT1 appeared to increase the nuclear export rate of *MYC* mRNA several-fold in comparison to siNC-treated cells, suggesting a link between *CCAT1* eRNA and *MYC* gating. We next explored whether the increased export rate correlated with the increased presence of *MYC* alleles at the nuclear periphery, typical of gated genes. DNA FISH analyses revealed that both the OSE, containing the *CCAT1* gene, and *MYC* alleles were indeed enriched more than twofold at the nuclear periphery in siCCAT1-treated cells compared to siNC control cells (**Fig. 1D**).

The significant inhibition of cell proliferation in siCCAT1-treated cells - reflected by spiked-in RNA recoveries (**Fig. S1A**) - raised the possibility that the increase in the nuclear export rate of *MYC* mRNA in this condition might, in part, be mediated by cell confluence-dependent mechanisms. In congruence with this supposition, RNA FISH analyses (**Fig. S1B**) revealed that active *MYC* alleles were significantly more frequent within 0.6 micrometers from the nuclear periphery^18^ upon either siCCAT1 treatment (**Fig. 1E**) or seeding the cells at 10%, 50% confluence compared to 100% confluence (**Fig. 1F**). To examine the relationship between cell confluence and mRNA export rates, we first determined the relationship between cell confluence as measured by IncuCyte and total RNA recovery, using spike-ins (**Fig. S1C**). Using RNA recovery as a proxy of cell confluence, the results revealed that the nuclear export rates of newly synthesized *MYC* mRNAs were indeed several-fold higher at low to medium cell confluence compared to confluent cells (**Fig. 1G**). Plotting the export data with the recoveries of corresponding samples documented that the nuclear export rates of *MYC* mRNA inversely correlate with cell confluence. Moreover, the nuclear export rates in si*CCAT1*-treated cells are similar to the export rates generated in cells at low confluence (**Fig. 1H**) despite the lower levels of *MYC* mRNA accumulation in these cells. We conclude that the gating of active *MYC* templates to nuclear pores is confluence-dependent. Moreover, the *CCAT1* eRNA, while activating *MYC* transcription, might indirectly counteract *MYC* gating, raising the possibility that *MYC* transcription itself impacted its gating frequency.

### MYC gating enables exponential cell growth concomitant with increased expression heterogeneity

To examine the relationships between *MYC* gating, transcription and cell confluence-dependent proliferation rate closer, we first compared the cell proliferation rate of wild type (WT) cells at different cell confluences to that of derived OSE-mutant cells (E3 and D4) that lack *MYC* gating but transcribe *MYC* at similar levels to WT cells despite reduced *CCAT1* expression^19^. **Figure S2A** shows that at cell confluences >50%, cell division of WT cells slowed significantly, likely reflecting partial contact inhibition. Following cell division every 3 h for a period of 72 h, the rate of cell proliferation (**Fig. S2B**) closely matched the nuclear export rates, as shown in **Figure 1H**. Next, WT, E4 and D3 mutant cells were seeded at low confluence, and the rate of cell proliferation was followed for 72 hours. **Figure S2C** shows that the mutant cells displayed a strongly reduced cell proliferation rate in comparison with the WT cells, suggesting that *MYC* gating is a major driver of exponential cell proliferation in colon cancer cells at low confluence and is itself regulated by cell confluence.

To assess the contribution of *MYC* gating to MYC levels and the potential influence of *MYC* transcription itself on the observed increase in *MYC* gating at low cell confluence, we first compared *MYC* mRNA accumulation to transcription at 10, 50 and 100% confluence. Since the half-life of *MYC* intron 1 is short (ca. 7 minutes) in cells harvested at an average cell confluence of 15.5% (**Fig. S2D**), labeling newly synthesized RNA with EU for 15 minutes is a comparatively slow process for estimating transcription, as it encompasses cellular uptake and conversion to ethynyl-UTP before incorporation into RNA. *MYC* transcription was therefore instead measured by quantitating *MYC* intron 1 levels in total RNA. Interestingly, *MYC* transcription rates inversely correlated with cell confluence, a pattern observed only in WT cells, indicating that a MYC-mediated control of *MYC* transcription^13^ requires a functional CTCF binding sites within the *CCAT1* gene distal to *MYC* (**Fig. 2A; Fig. S2E**). It is currently unclear why *MYC* transcription rate increased in the mutant cells at low confluence, as they largely lack *CCAT1* expression^19^. One possibility is that other neighboring, non-coding genes, such as *CASC8*^*21*^, which is overexpressed in E4 cells (preliminary results), compensates for the loss of *CCAT1* expression. To more precisely assess the contribution of *MYC* gating to the increasing *MYC* mRNA accumulation in WT cells at low cell confluence, we employed our published algorithm^18^ to simulate *MYC* mRNA accumulation using the values of transcription levels, *MYC* mRNA nuclear export rates and nuclear and cytoplasmic *MYC* mRNA degradation rates from **Figures 1C, G; Figures S2D-F**. Setting the nuclear export rates at 8 or 1 (the latter representing a nongated condition^19^) closely recapitulated the observed *MYC* mRNA accumulation values (**Fig. 2B**). We conclude that the frequency of *MYC* gating is a major determinant of the total level of *MYC* mRNA expression in low-to mid-confluent cells, paralleling reduced transcriptional rates.

**Figure 2.**
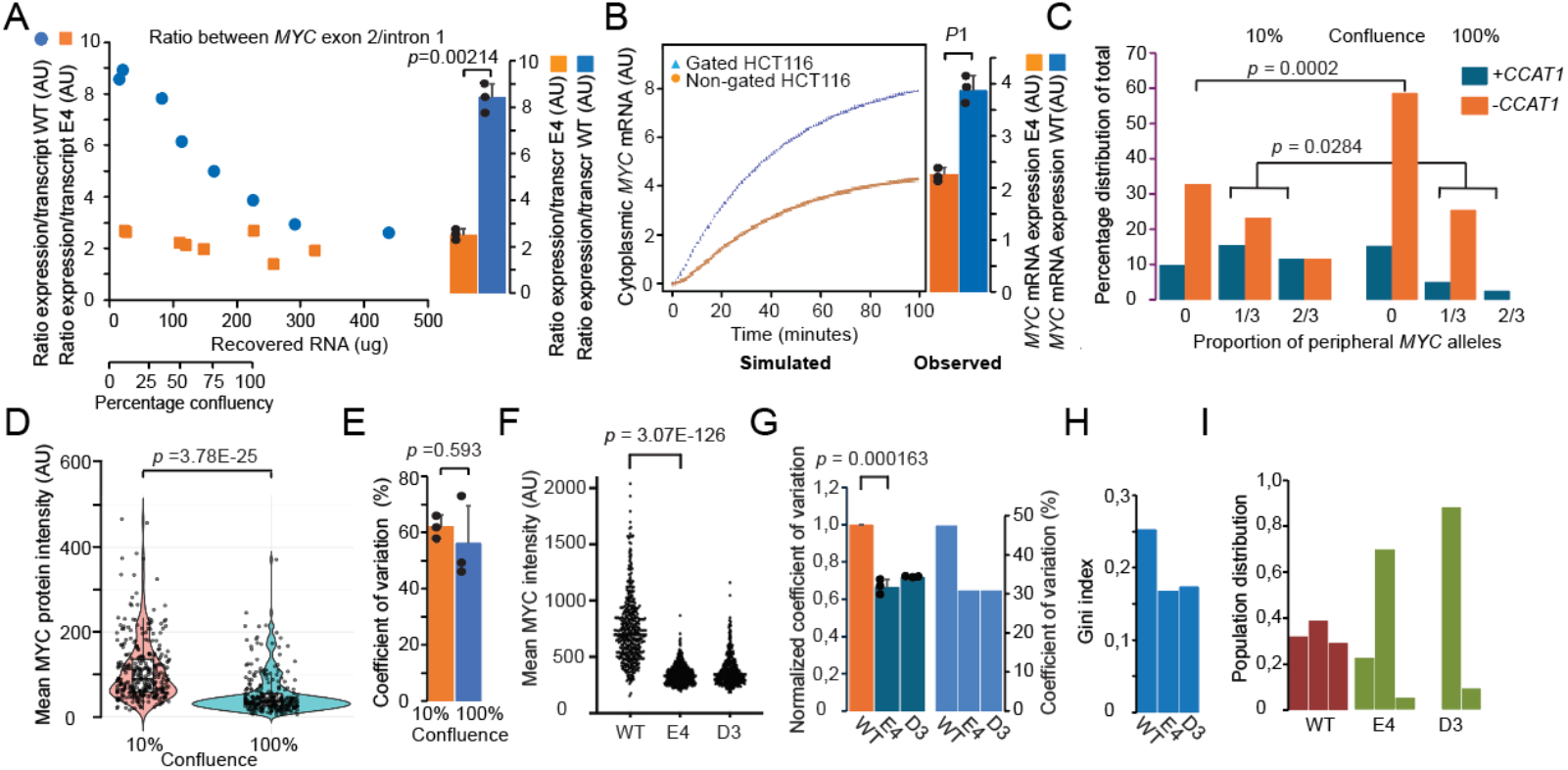
MYC gating drives both exponential cell proliferation and MYC expression heterogeneity. A) Ratios between total *MYC* mRNA accumulation and transcription (from **Fig S2E-F**) to visualize the output of mRNAs in relation to transcriptional activity. B) Simulation of the contribution of gating to total *MYC* mRNA accumulation (right, diagram) by integrating mRNA decay data, nuclear export rates and transcription rates, as previously described^18,19^. The blue and red lines depict the values obtained via nuclear export rates of 8 (gated) and 1 (nongated), respectively. P1 = 8.0897E-05 C) RNA FISH analyses of simultaneous transcription of *MYC* and *CCAT1* in individual cells at different confluences. Violin plot (D) and coefficient of variation (CV; E) analyses of mean MYC expression in HCT116 cells at 10 (199) or 100% (224) confluence. The data is the sum of three independent experiments. F) Scatter plot analyses of mean MYC expression of WT and mutant (D3, E4) cells with MYC expression plasticity analyzed by CV (G), Gini index (H), and Mclust distribution (I) analyses. A total of 1443 cells were analyzed. The *P* values were determined by two-tailed unpaired Student’s t test (A,B,D-G) or Fisher’s exact test (C).

Given that the CTCF binding site within the OSE positively regulates both *MYC* gating^19^ and *CCAT1* expression - which antagonizes *MYC* gating - in a sub-nuclear compartment-dependent manner, we hypothesized that the coordination of these processes might result into increased cell-to-cell heterogeneity of MYC expression even at high mean MYC levels. Peripheral active *MYC* alleles thus often co-occur with non-peripheral, active *CCAT1* alleles in the same nuclei by RNA FISH analyses, suggesting dynamic transitions between gated and non-gated expression patterns in response to cell confluence (**Fig. 2C**). Moreover, the same cell can display both peripheral and intra-nuclear *MYC* alleles with inactive and active *CCAT1* alleles, respectively, suggesting that the *CCAT1* eRNA acts mainly locally. Using optimized conditions for nuclear MYC immunostaining (**Fig. S2G**), we could document that the average level of mean MYC protein expression per cell increased ca 2-fold from high to low cell confluence (**Fig. 2D**), without decreasing its coefficient of variation (CV, **Fig. 2E**). This was a surprising result as increased MYC expression during exponential growth would be expected to reduce expression heterogeneity. This trend was reinforced by the observation that despite a >2-fold higher mean MYC expression level in WT than the mutant cells (**Fig. 2F**) and similar *MYC* transcription rates in all the three cell lines^19^, WT cells displayed the highest CV value of mean nuclear MYC expression relative to both the slow-growing E4 and D3 cells (**Fig. 2G**). Global variance tests (Fligner–Killeen and Levene) confirmed these highly significant differences in dispersion across genotypes (p < 10^−50^). Information-theoretic measures provided an orthogonal view of this heterogeneity, with WT cells exhibiting the greatest Gini index (**Fig. 2H**), indicating a broad and unequal distribution of MYC expression levels across the cell population. In contrast, the E4 and D3 mutants presented reduced inequality, which is consistent with more uniform expression. Importantly, Gaussian mixture modeling further revealed that MYC expression in WT cells is best described by three overlapping subcomponents, whereas E4 and D3 cells are dominated by a single major component, with only minor high-MYC subfractions (**Fig. 2I**). Dip test p values (> 0.3) indicated that these subpopulations form a continuum rather than discrete peaks. We conclude that genetic evidence supports *MYC* gating as one key factor in simultaneously increasing both the amplitude of MYC expression heterogeneity and mean MYC protein levels.

### MYC transcriptional rate regulates MYC gating

As **Figure 2A** hinted at a role of transcription rates in controlling the frequency of *MYC* gating, we analyzed the volumes of RNA FISH signals (identified by sequential RNA and DNA FISH analyses). The results show that weak RNA FISH signals, potentially identifying transcriptionally paused *MYC* alleles, appeared to accumulate at the nuclear periphery especially at low cell confluence (**Fig. 3A**). The link between the transcription rate and the gating process was further supported by the observation that the silencing of *CCAT1* expression significantly reduced the peak intensity of *MYC* intron and exon probes and the volume of *MYC* exon probes (**Fig. 3B**), indicating a marked reduction in the transcriptional initiation and/or elongation rate. As Ser5 phosphorylation of Pol II marks transcriptional initiation at active promoters and gradually decreases over the gene body during transcriptional elongation^22^, we first determined the role of *CCAT1* eRNA in this process. Whereas the patterns generated in siNC-treated cells were, as expected, gradually reduced along the gene, the level of Ser5-phosphorylated Pol II was reduced in the 5’of the *MYC* gene and remained constantly low in siCCAT1-treated cells (**Fig. 3C**). Moreover, the ratios of total Pol II occupancy between siCCAT1- and siNC-treated cells increased significantly along *MYC* to indicate reduced elongation rate^23^ in addition to a reduction in the re-initiation of *MYC* transcription in the relative absence of *CCAT1* eRNA (**Fig. 3D**). As the levels of total pol II are reduced 5-fold at the first exon in siCCAT1-treated cells, it appears that the pausing of the elongation rate is mechanistically different from pausing at the promoter-proximal region^23^. We conclude that the *CCAT1* eRNA function promotes both transcriptional initiation and elongation of *MYC* while antagonizing the nuclear export rate of *MYC* mRNAs. Interestingly, the *CCAT1* gene (positioned within the OSE) in contrast to *MYC* reaches the nuclear periphery/pore in an inactive state (**Fig. 1D, E**), suggesting that absence of local *CCAT1* eRNA triggers the migration of OSE-*MYC* complex to nuclear pores.

**Figure 3.**
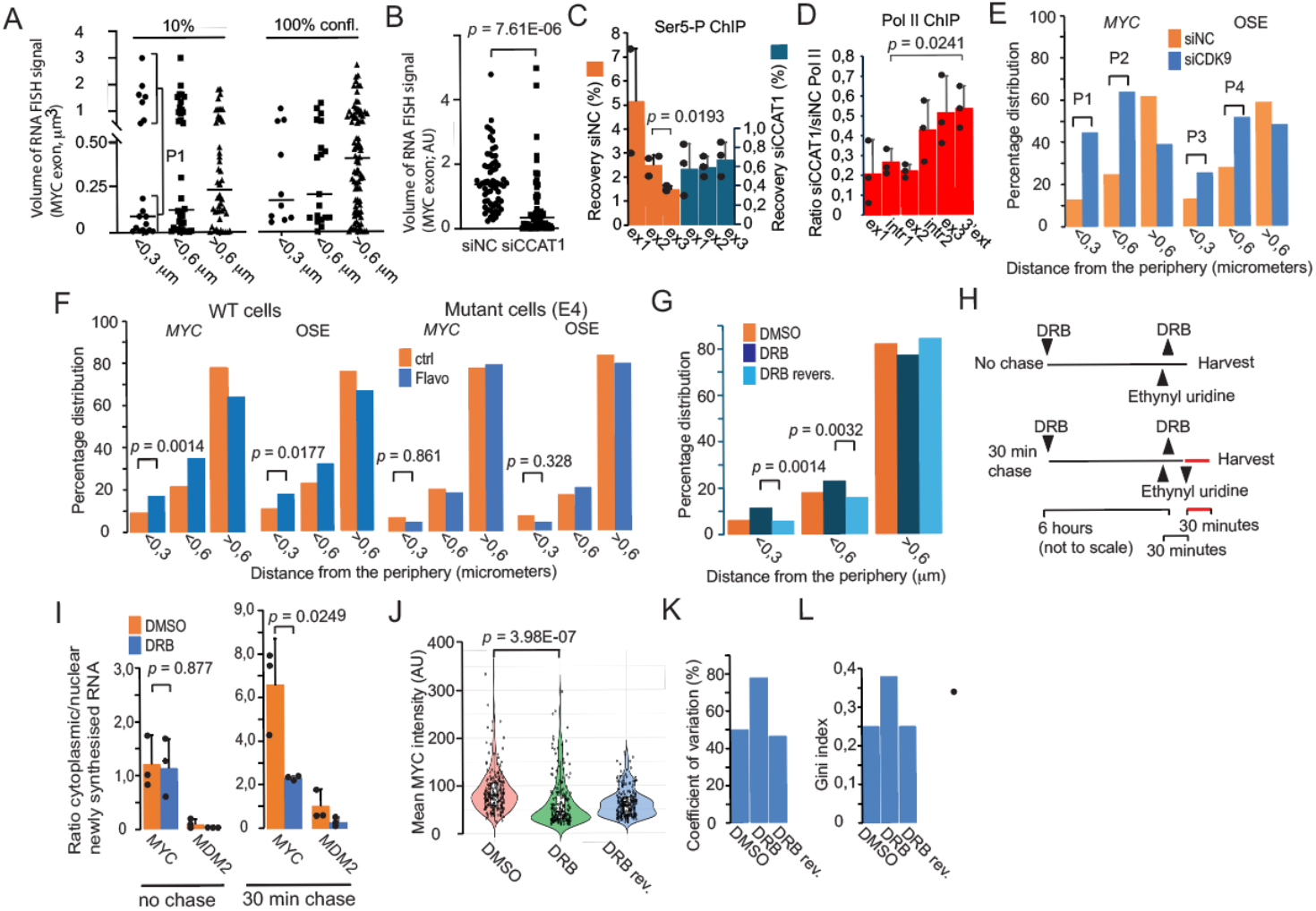
Transcriptional rate regulates MYC gating. A) The volume of RNA FISH signals, marked by *MYC* exon probe in cells grown at 10% (49 alleles) or 100% (88 alleles) confluence. The nuclear *MYC* signals were identified by sequential RNA and DNA FISH analyses. P1 = 9.32E-06. B) Volumes of RNA FISH signals in control and siCCAT1-treated cells based on the counting of 65 (*MYC*, siNC) and 57 (*MYC*, siCCAT1) active alleles. ChIP analyses of the distribution of Ser-5-phosphorylated (C) or total (D) Pol II along the *MYC* gene in siNC- and siCCAT1-treated cells. Ex1 = exon 1; intr1 = intron 1; ext = 3’ of *MYC* (**Fig. S3A**). E) WT HCT116 cells were transfected with siCDK9 or siNC, followed by analyses of the distribution of *MYC* alleles in relation to the nuclear periphery *via* DNA FISH. A total of 152 alleles were counted for both siNC- and siCDK9-treated cells and represent the average of two independent experiments with a siCDK9 knockdown efficiency of >80% in both cases. The cells were seeded at50% confluency. P1 = <0.0001; P2 = <0.0001; P3 = 0.0025; P4 = 0.0027. F) Flavopiridol treatment increases the presence of *MYC* and OSE alleles at the nuclear periphery in WT cells (left) but not in mutant E4 cells (right). The results are the average of two independent experiments with a total of 379 (WT) and 270 (E4) alleles counted. G) DRB treatment increases the presence of active *MYC* alleles at the nuclear periphery in WT HCT116 cells. The RNA FISH analyses, which used a probe for the entire *MYC* gene **(Fig. S3B**), represent the average of three independent experiments (fixed at50% confluence), with a total of 424 (DMSO), 460 (DRB) and 591 (DRB reversal) alleles counted. H) Scheme for exploring the effects of reversing the effects of DRB treatment on nuclear export rates. I) Nuclear export rates of mRNAs derived from *MYC* and *MDM2* following DRB reversal with no chase (left image) or 30 minutes of chase (right image). The cells were seeded at 50% confluence. J) Violin scatter plots showing the distribution of the mean nuclear MYC protein signals in HCT116 cells treated with DMSO, DRB, or DRB reversal, as above. Heterogeneity metrics of CV (K) and Gini index (L) analyses of the cell-to-cell heterogeneity in mean MYC protein levels. The results are the sum of three independent experiments unless otherwise indicated. The *P* values were determined *via* unpaired, two-tailed Student’s t tests (A-D, I, J) or Fisher’s exact tests (E-G).

In the absence of robust *in situ* techniques to examine the spatial distribution of transcriptional elongation rates in the 3D nuclear architecture, we exploited our previous observation that the inhibition of transcriptional elongation by Flavopiridol drove circadian genes to the nuclear periphery^24^. Indeed, siCDK9 (**Fig. 3E**) and Flavopiridol (**Fig. 3F**) treatments similarly increased the presence of both the OSE and *MYC* alleles at the nuclear periphery. Notably, the Flavopiridol-dependent effect was absent in E4 cells carrying a mutated CTCF binding site within the *CCAT1* intron^19^ (**Fig. 3F**), indicating that the CTCF binding site within the distal OSE is essential for the recruitment of paused *MYC* alleles to the nuclear periphery. Another drug, DRB, that targets the elongation complex has the combined benefit of both inhibiting the elongation process and being rapidly reversible^25^ by selectively targeting Pol II at regions extending up to a few kb from the promoter^26^. Accordingly, DRB promoted the juxtaposition of both OSE and *MYC* alleles to the nuclear periphery (**Fig. S3A**), with an increased frequency of *MYC* RNA FISH signals juxtaposed to the nuclear periphery after 6 hours of DRB treatment (**Fig. 3G, Fig. S3B)**. Intriguingly, a 20-minute wash out of DRB (DRB reversal, **Fig. 3G**) was sufficient to reduce this effect, suggesting that the template is released from the nuclear pore once transcription is completed.

To examine whether the DRB-induced migration of active *MYC* alleles to the nuclear periphery would enable the quick recovery of nuclear export rates upon DRB reversal, RNA was ethynyl-uridine (EU)-labeled for 30 minutes starting 10 minutes before the end of the DRB treatment, allowing for uptake and conversion to EU-UTP, followed by another 20 minutes of labeling after the reversal of DRB (**Fig. 3H**). The cells, seeded at midconfluence, were either harvested directly after EU labeling (0-hour chase) or chased for 30 minutes before harvesting. Strikingly, the nuclear export rate of *MYC* mRNA was indeed similar in the 0-hour samples and control samples, and it increased slightly at the 30-minute chase time point compared to the 0-hour time point of DRB-pretreated cells (**Fig. 3I**). In contrast, there was no detectable presence of newly synthesized cytoplasmic mRNA derived from the nongated *MDM2* in the 0-hour DRB-pretreated sample (**Fig. 3I**) despite a similar degree of transcriptional reactivation **(Fig. S3C**). We argue, therefore, that paused peripheral *MYC* alleles rapidly complete their transcription at nuclear pores upon removal of DRB, followed by the facilitated export of derived mRNAs and the release of the *MYC* gene from the nuclear pores/periphery into the nuclear interior. Importantly, Violin scatter plots of the DMSO control and DRB-treated cells indeed revealed that the DRB-treatment generated significant changes in MYC protein expression levels (**Fig. 3J**) with increasing cell-to-cell heterogeneity, as determined by the CVs (**Fig. 3K**) and the Gini indices (**Fig. 3L**). These observations indicated that DRB treatment induced a broader and more unequal distribution of MYC protein levels, and these features were reduced upon DRB reversal. Whereas global variance tests indicated significant differences across the three conditions (Fligner *p* ≈ 0.016; Levene *p* ≈ 0.0023), Hartigan’s dip test did not support multimodal distributions in any condition (all dip p > 0.55). These data revealed that DRB treatment reduced the mean nuclear MYC intensity while increasing its relative heterogeneity and inequality in the cell population. Following drug reversal and the release of the *MYC* gene from the nuclear periphery, MYC expression levels remained low, but their variance decreased significantly, returning to or below normal levels in control cells.

### The MYC protein drives MYC gating in a positive feedback loop

Given the early observations that MYC regulates the transcription of its own template in a negative feedback loop^3^, we reasoned that high MYC expression levels should have a positive effect on gated expression. Accordingly, knockdown of *MYC* expression increased *MYC* transcription approximately 3-fold (**Fig. 4A**) with *MYC* transcripts diffusely distributed within the nucleus (**Fig. 4B**), suggesting impaired export. In congruence with the observations above, the reduction of MYC expression increased *CCAT1* expression (**Fig. 4A**), potentially explaining the increased transcriptional rate of *MYC* in siMYC-treated cells. We note that this result contradicts an earlier observation arguing that MYC activates *CCAT1*^*1*6^, which could reflect different experimental conditions. Accordingly, nuclear export assays revealed that the nuclear export rate of *MYC* mRNAs in si*MYC*-treated cells were reduced to the level observed in nongating mutant cells—while the export rates of the nongated *MDM2* and *AHCTF1* genes were not affected by the absence of MYC (**Fig. 4C**). This effect, which was prominent at low cell confluence (**Fig. 4D**), partially reflected a significant reduction in the recruitment of active *MYC* alleles to the nuclear periphery in siMYC-treated cells (**Fig. 4E**). This observation agrees, moreover, with the notion above that high *MYC* transcription rate anticorrelates with hallmarks of *MYC* gating. We conclude that the MYC protein promotes the recruitment of active *MYC* alleles to the nuclear periphery and increases the nuclear export rate of *MYC* mRNAs to increase *MYC* mRNA accumulation despite reduced *MYC* transcription. This positive feedback loop is likely driven by the negative feedback of MYC on its own transcription, potentially mediated by reduced *CCAT1* expression.

**Figure 4.**
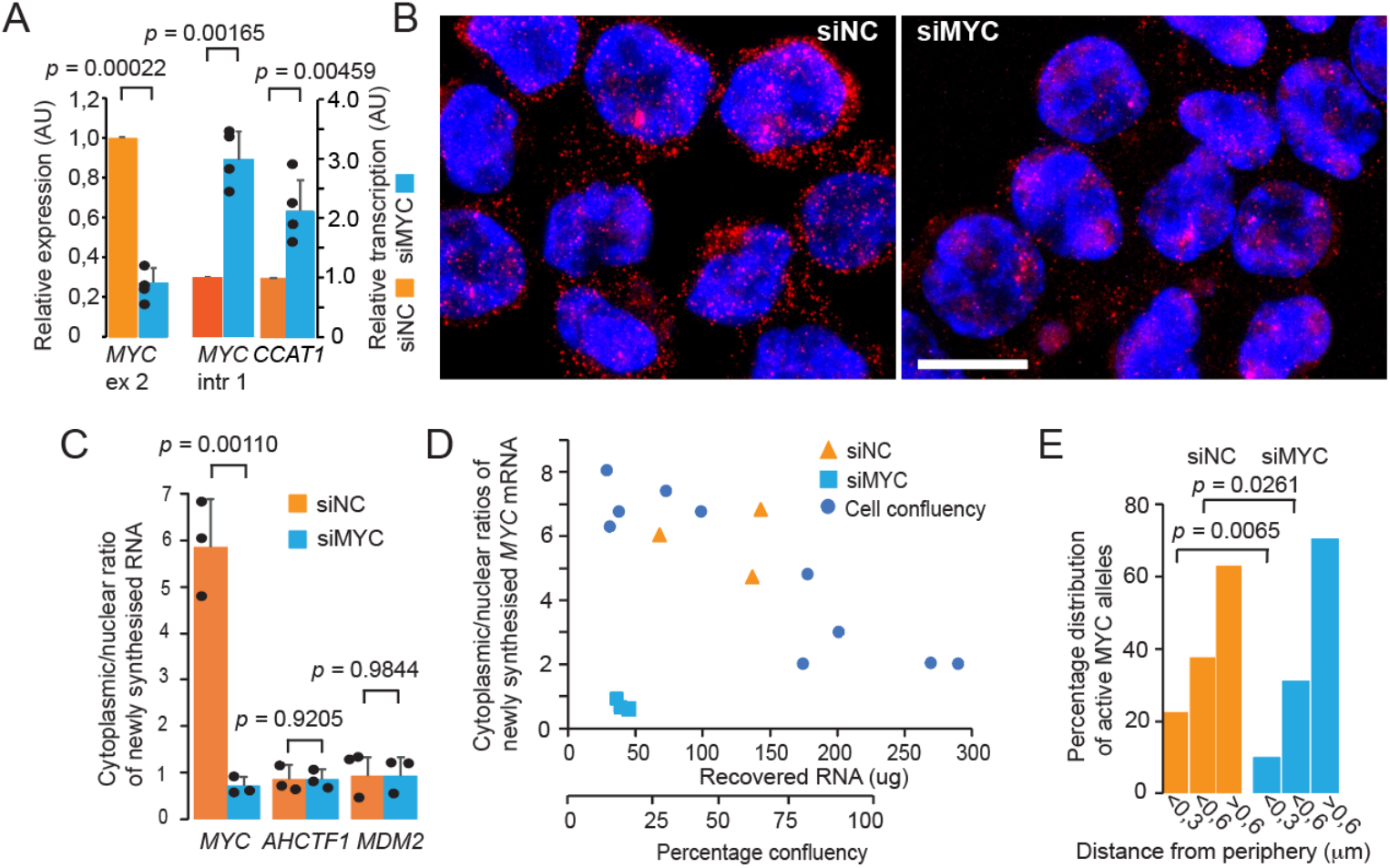
MYC expression drives MYC gating. RT-qPCR analyses of *MYC* exon 2 (left) and intron 1 (middle) and *CCAT1* exon 2 (right) in total RNA isolated from siNC- or siMYC-treated cells. The downregulation of *MYC* mRNA in siMYC-treated cells was >70% on average in four independent experiments. B) Representative *MYC* RNA FISH images using Stellaris MYC exon probes in siNC- or siMYC-treated cells. C) Nuclear export rates of *MYC* mRNA in control (siNC)- and siMYC-treated cells seeded at low confluence, represented by three independent experiments. Two nongated genes, *AHCTF1* and *MDM2*, served as internal controls, indicating that siMYC treatment did not affect the nuclear export rates of their derived mRNAs. D) The nuclear export rates were related to cell confluence, as determined by the recovery of total RNA. Notably, the siMYC-treated cells represent low confluence, likely due to the inhibition of cell cycle progression. E) Distribution of active *MYC* alleles in relation to the nuclear periphery in siNC- or siMYC-treated cells. The data were generated from two independent experiments with a knockdown efficiency of *MYC* expression >90% in both cases. A total of 131 (siNC) and 121 (siMYC) active *MYC* alleles were counted. The *P* values were determined by one-sample (A), two-tailed unpaired Student’s t test (C) or Fisher’s exact test (E). Bar = 10 μm.

Figure 5 schematically summarizes these observations, outlining how one negative feedback loop promotes another, positive feedback loop to generate both high and variable MYC expression. As a driver of *MYC* transcriptional initiation and elongation, *CCAT1* eRNA counteracts gated MYC expression. In line with an antagonistic role of *CCAT1* eRNA to *MYC* gating, its gene is mostly inactive at the nuclear periphery. This scenario identifies the silencing of *CCAT1* and hence reduction of *MYC* transcriptional activity as a potential trigger for the migration of the paused *MYC* to the nuclear pores. We also envisage a scenario where extrinsic cues, such as WNT, which activate both *CCAT1* transcription^19^ and *MYC* gating^19^ promote cycles of interdependent *CCAT1* and *MYC* expression events while continuously antagonizing each other. Due to the unique relationship between the *CCAT1* function and *MYC* gating, increased mean MYC expression is accompanied by increased cell-to-cell heterogeneity, opposite to what would be expected at high expression levels.

**Figure 5.**
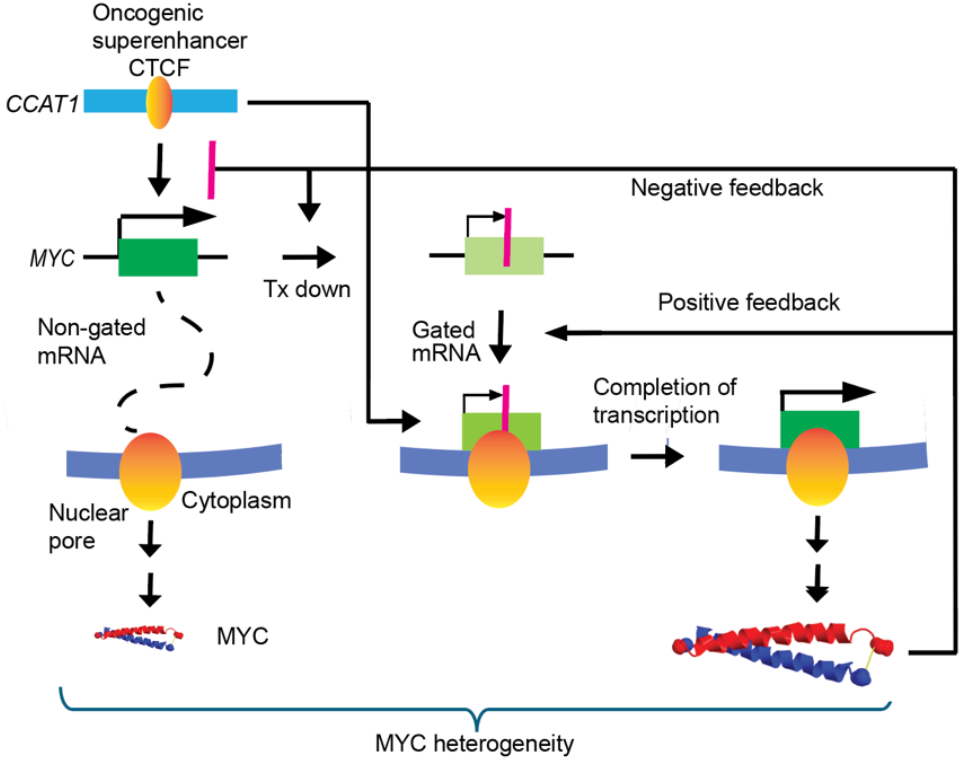
A schematic summary of the negative feedforward principles underlying *MYC* gating induced MYC expression heterogeneity. Positive and negative feedback loops indicated in the image are likely required to manifest the gating process. We envisage in the scheme that pathological expression of the MYC protein reduces *CCAT1* expression, which indirectly promotes gated expression by attenuating *MYC* transcription. As the initiation of transcription promoted by *CCAT1* eRNAs is a likely prerequisite for the subsequent gating process, we submit that transient reductions on transcription rates will trigger the migration of *MYC* to the nuclear pore. This scheme assumes that transcriptional rates are heterogeneous and subject to the effects of intermittent extrinsic cues to generate expression noise that is manifested by the balance between gated and nongated MYC expression. As the CTCF binding site within the oncogenic super-enhancer directs both *CCAT1* expression and *MYC* gating, it constitutes a master switch in this scheme.

### Hallmarks of MYC gating in invasive colorectal and breast tumors correlate with expression plasticity

To assess to what extent these observations extend from a model system to the *in vivo* situation, we first confirmed that peripheral *MYC* distribution, a hallmark of *MYC* gating, is a predictor of an increased nuclear export rate of *MYC* mRNA in HCT116 cells (**Fig. 6A**). Next, we addressed the presence of *MYC* gating hallmarks in a primary luminal A breast cancer sample stained for NUP133, with selected regions marked with different color codes (**Fig. 6B**). Quantitation of the subnuclear localization of the *MYC* DNA FISH signals revealed a very high frequency of their peripheral (<0.6 micrometers from the periphery) distribution in both the Ductal Carcinoma *In Situ* (DCIS) (blue) regions within the tumor and invasive cells (red/orange boxes; **Fig. 6B, C**). Chromatin *in situ* proximity (ChrISP) analysis^18,19,27,28^ revealed close physical proximity between OSE DNA FISH probes and NUP133 epitopes at the nuclear periphery of DCIS cells (**Fig. 6D**, marked with a white box in panel B). Clusters of *MYC* copies were, moreover, in close physical proximity (<0,6 micrometers) to the nuclear periphery in both metastatic and primary CRCs (**Fig. S6A**). The quantitative difference in patterns of *MYC* distribution was particularly striking in comparison with primary cultures of human colon epithelial cells with *MYC* alleles almost completely depleted from the nuclear periphery (**Fig. S6B**).

**Figure 6.**
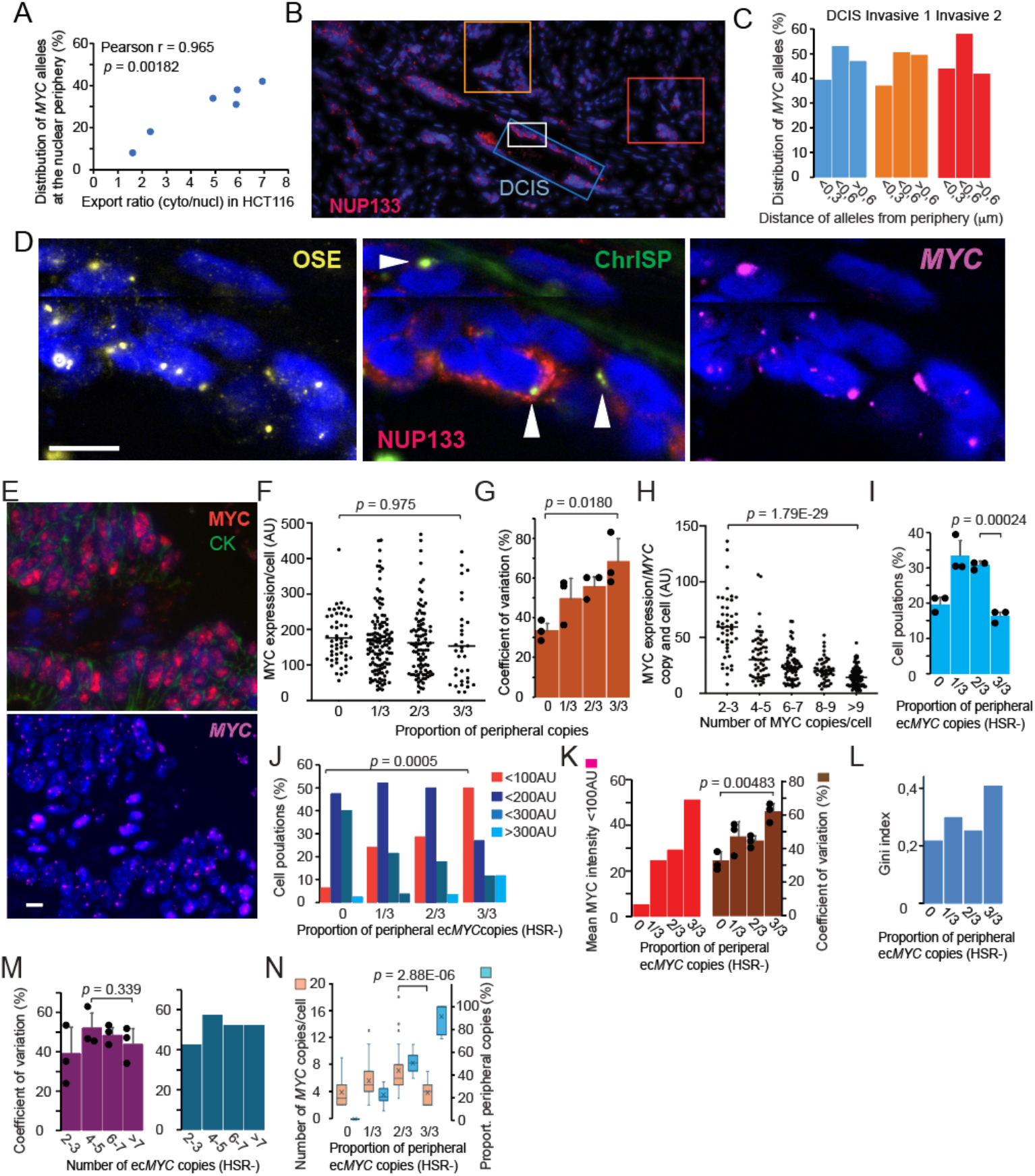
Hallmarks of MYC gating in primary and metastatic tumors correlate with MYC expression plasticity. A) Proximities (within 0.6 μm) of *MYC* alleles to the nuclear periphery predict the nuclear export rates of *MYC* mRNAs in HCT116 cells. B) Overview of an invasive breast cancer tissue containing a *ductal carcinoma in situ* (DCIS) region strongly stained for NUP133 expression. The color-coded, boxed areas identify regions selected for quantitation of the spatial distribution of *MYC* alleles (C; 263 *MYC* alleles were counted). The white box of panel B is enlarged in panel D to illustrate ChrISP signals (arrows), generated between the super-enhancer (OSE) sequences and NUP133 when in sufficient proximity (<160 Å) to each other^18^. E) Representative view of MYC protein immunofluorescence signals combined with *MYC* DNA FISH signals. CK = cytokeratin. F) Scatter plots of mean nuclear MYC protein expression levels in relation to the proportion of peripheral *MYC* copies/cell. The coefficient of variation of the mean intensity of nuclear MYC staining was determined on a cell-to-cell basis for the three tumors in relation to the proportion of peripheral *MYC* copies (G). H) MYC protein output per cell per *MYC* copy. The data are based on a total of 304 cells (1242 *MYC* alleles). I) Distribution of HSR-negative cells from three different CRC samples stratified by the proportion of peripheral ec*MYC* copies. J) Bar diagram comparing MYC expression levels in HSR-negative cell populations (color-coded in the image) stratified by the proportion of peripheral ec*MYC* copies. K) The prominent (68–100%) peripheral distribution of ec*MYC* copies (left) is correlated with an increased coefficient of variation in the mean nuclear protein MYC levels (right). L) Gini index of MYC expression variability in relation to the proportion of peripheral ec*MYC* copies in HSR tumor cells. M) Coefficient of variation of mean nuclear MYC levels in cells stratified by the total number of ec*MYC* copies (excluding HSR*MYC*) in individual tumors (left) and when combined (right). N) Box plot analyses comparing the distribution of ec*MYC* copy numbers with the proportion (proport.) of peripheral ec*MYC* copies. The *P* values were determined by Pearson (A), the unpaired, two-tailed Student’s t test (F-I, K-M) and Fisher’s exact test (J). The bars in (D) and (E) corresponds to 10 μm.

Next, to explore if the spatial distribution of *MYC* correlated with expression heterogeneity, we subjected three primary CRC samples to a combination of MYC protein immunostaining and DNA FISH-based quantitation of the peripheral distribution of *MYC* (**Fig. 6E**). Visualization of the distribution of the mean nuclear MYC protein levels in four categories of cells with increasing proportions of peripheral *MYC* copies revealed a gradual increase in the cell-to-cell heterogeneity of the mean nuclear MYC levels (**Fig. 6F**). The CV values of the mean nuclear MYC levels thus correlated with the proportion of peripheral *MYC* copies in each individual CRC sample, increasing more than twofold at the highest proportion of peripheral *MYC* copies (**Fig. 6G**). Intriguingly, the highest CV values correlated with only an average of 4 *MYC* copies at the periphery (**Fig. 6G; Fig. S6C**), indicating that the *MYC* copy number did not *per se* translate into MYC expression variation in these tumor samples. Accordingly, the CV values showed only a weak and statistically insignificant increase when cells were sorted according to the number of *MYC* copies, without taking the peripheral distribution of *MYC* into account (**Fig. S6D**). Notably, the relative output of MYC protein signals per *MYC* copy seemed to decrease with increasing copy number (**Fig. 6H**), which is consistent with an active, negative feedback of MYC on its transcription in all three CRC samples.

The simultaneous presence of ec*MYC* and *MYC* HSRs (HSR*MYC*) in a subpopulation of tumor cells (**Fig. S6E, F**) complicated, however, the relationship between *MYC* copy numbers and MYC expression profiles. Robust technical approaches to precisely assess *MYC* copy numbers in HSRs *in situ* do not yet exist to hamper a direct comparison between HSRs and MYC expression plasticity. We therefore first assessed whether the presence of HSR*MYC* affected the spatial distribution of ec*MYC* copies by stratifying the cells into HSR- and HSR+ cell populations. This approach revealed that in the presence of HSR*MYC*, the proportion of peripheral ec*MYC* copies was moderately increased (**Fig. S6G**), and the correlation between the mean MYC CV and the peripheral localization of *MYC* copies (ecDNA+HSRs) was attenuated without changing overall CVs in the tumors (**Fig. S6H**). These observations revealed that ec*MYC* copies are sufficient for generating heterogeneous MYC expression. Notably, while the mean MYC expression levels decreased in proportion to the number of peripheral ec*MYC* copies, it increased with reduced CVs in cells containing a combination of ec*MYC* and HSR*MYC* copies (**Fig. S6I**), suggesting that HSR*MYC* copies antagonize the effect of ec*MYC* copies on expression heterogeneity while increasing mean MYC levels.

Based on observations in HCT116 cells, it could be expected that the population densities of tumor cell populations would be redistributed according to subnuclear localization of ec*MYC* copies, linked with MYC expression variation. While the overall cell population distributions were similar between the tumors (**Fig. 6J**), violin and scatter plots indeed revealed that these were re-distributed according to the proportion of peripheral ec*MYC* copies and the mean level of MYC expression signals (**Fig. S6J, K**). Moreover, an increasing proportion of peripheral ec*MYC* copies correlated with a gradual reduction in the overall mean MYC signal (**Fig. S6I-K**). The clustering of low-MYC cells (<100 AU, red box in **Figure S6J**) with peripheral ec*MYC* copies (**Fig. 6J**) correlated with an overall increase in CVs (**Fig. 6K**) and Gini indices of mean cellular MYC levels (**Fig. 6L**). Care was taken to ensure that the low-MYC cells corresponded to tumor cells within the tumor mass and were marked with cytokeratin and *MYC* amplification (**Fig. S6L**). The absence of a link between CV values of mean cellular MYC levels and ec*MYC* copy numbers *per se* in HSR-cells (**Fig. 6M**) reinforces the correlation between ec*MYC* gating and increased MYC expression variation. Finally, the data suggest that the heterogeneity of the mean MYC signals was optimized at 3–5 *MYC* copies per cell in these tumors (**Fig. 6N**), which is in keeping with that the costs of maintaining too many ec*MYC* copies anticorrelates with cancer cell plasticity^8^.

## Discussion

We have shown here that *MYC* gating to nuclear pores is a key driver of the exponential division of cancer cells, a process balanced by the influence of the *CCAT1* eRNA on the transcriptional activity of *MYC*—the greater the transcription, the less prominent the gating and *vice versa*. Accordingly, high MYC protein expression limits *MYC* transcription while concomitantly increasing the *MYC* gating frequency in a positive feedback loop. Importantly, genetic evidence shows that *MYC* gating paradoxically also increases MYC protein cell–to-cell heterogeneity, as shown by heterogeneity metrics. *MYC* gating in combination with its antagonist, the *CCAT1* eRNA, amplifies transcriptional and gene expression diversity within the cell population; without it, tighter, more homogeneous MYC regulation emerges to potentially limit phenotypic plasticity.

Several lines of evidence point to a regulatory role of transcriptional activity in the *MYC* gating process. First, the nuclear export rate of *MYC* mRNAs in si*CCAT1*-treated cells increased according to cell confluence, although the expression of MYC was reduced to approximately 50% in siCCAT1-treated cells. Second, the volumes of the RNA FISH signals were strongly reduced in siCCAT1-treated cells, indicating that a reduction in transcriptional activity paralleled the increase in *MYC* mRNA nuclear export rates. Third, the nuclear export rate of *MYC* mRNAs inversely correlated with *MYC* transcriptional activity along a spectrum of cell confluence from low to high. Fourth, knocking down overall MYC expression increased *MYC* transcription while antagonizing the facilitated nuclear export of *MYC* mRNA. Fifth, the elongation inhibitors Flavopiridol, siCDK9 and DRB facilitated the recruitment of *MYC* alleles to the nuclear periphery. Importantly, the active *MYC* alleles retain, to a large degree, their template-associated transcripts at the nuclear periphery during 6 hours of DRB treatment. Upon DRB reversal, the nuclear export rates are very similar to those of the control cells at 0 hours, which likely represents the completion of paused transcription at *MYC* alleles initially driven to the nuclear periphery by DRB treatment. Sixth, this interpretation aligns with the biphasic transcriptional activity of peripheral *MYC* alleles in low-confluency cells, suggesting that the *MYC* alleles reach the nuclear pore in a paused state, followed by the completion of transcription in this nuclear sub-compartment. Taken together, these observations document that the rate of *MYC* transcription is a regulator of *MYC* gating in HCT116 cells.

The influence of *MYC* gating on MYC expression heterogeneity constitutes a central and paradoxical message of this report, as *MYC* gating also drives exponential cell growth. Thus, both genetic data in the model system and correlative data in the tumor samples support a role for *MYC* gating in increasing MYC expression heterogeneity. Importantly, as the effect is most pronounced in low confluent HCT116 cells, the question arises as to how this feature is controlled in tumor cells which continuously battle cell contact inhibition. One possibility is that *MYC* gating in the tumor microenvironment is dependent on signals, such as WNT^19^ and the Hippo pathway^29^, and mutations that antagonize contact inhibition-mediated effects. Such processes may impinge on the CTCF-dependent *MYC* gating process by simultaneously promoting both agonistic (AHCTF1-mediated recruitment to nuclear pores^19^) and antagonistic (*CCAT1* eRNA expression) downstream events.

The loss of active MYC templates from the periphery upon DRB treatment reversal indicates that once transcription has been completed at the nuclear pore, the templates are released into the nucleoplasm. The dynamics of ec*MYC* release from the nuclear periphery might influence the emergence of tumor cells with a very high proportion of peripheral *MYC* copies but simultaneously reduced levels of mean MYC protein expression. It is probable that *MYC* copies in such cells respond to the dynamic lack of extrinsic cues by reduced transcription rate and consequent recruitment to the nuclear periphery, where they might remain until a new wave of cues enables completion of transcription. It might therefore be important to consider that tumor cells with a high proportion of peripheral *MYC* copies could survive therapeutic interventions targeting WNT signaling, for example, to create opportunities for the cancer to relapse once the drug levels decrease. By analogy to the DRB experiments, such peripheral ec*MYC* copies might render cancer cells more sensitive to subsequent signaling cues and underlie gating-dependent MYC expression plasticity. The negative feedforward loop identified here may complicate targeting MYC expression in therapy regiments, especially as compensatory mechanisms might be activated during treatments, as indicated here. However, as the cells displaying peripheral *MYC* copies also display highest MYC expression plasticity, we identify the *MYC* gating process as a new opportunity to target tumor cell plasticity and hence therapy resistance.

## Limitations of the study

The approach of inferring dynamics by taking snapshots of ongoing events carries the risk of misrepresenting a process as being linear when it might in fact be non-linear. To deal with this issue, we have frequently used several approaches to examine the “snapshot dynamics” from different angles. This is exemplified by basing our argument that *MYC* transcriptional rate regulates the gating of its template on three different approaches: i) confluency parameters, ii) knocking down either *CCAT1* or *MYC* expression, iii) ChIP analyses of Pol II distributions and iv) using agents to inhibit elongation rates. Another limitation concerns the tumor data, which are entirely descriptive based on DNA FISH and MYC expression parameters, as RNA in thin tissue sections is degraded due to clinical practices. Nonetheless, we consider the data important as they provide an *in vivo* corroboration of data generated in an *in vitro* cell model system.

## Star Methods

### Study participants

The primary CRC samples were obtained at Akershus University Hospital (protocol approval by the Regional Committee for Medical and Health Research Ethics of Southeast Norway ID 350208 and the Institutional Review Board). The CRC liver metastatic tumor was obtained from a male patient in his 70s at Karolinska University Hospital after surgical resection with curative intent (approval 2020-05598, The Swedish Ethical Review Authority). The luminal A breast tumor material was collected at Karolinska University Hospital after anonymization; therefore, no ethical approval was needed in Sweden.

All tumor samples were selected randomly and collected after surgical resection with curative intent without neoadjuvant chemotherapy. Tumor 042 is a rectal surgical specimen from a female patient (aged 64 years) classified as pT3N0 with a mutation in KRAS (G12D) but identified as wild type for BRAF and NRAS. Tumor 069 is a surgical specimen from the sigmoid colon of a male patient (aged 48 years), classified as pT4N0 with mutations in KRAS (G12S) and identified as wild type for BRAF and NRAS. Tumor 078 is also a surgical specimen from the sigmoid colon of a male patient (aged 47 years) and is classified as pT4aN1b, with wild-type sequences in KRAS, BRAF and NRAS. All three tumors were microsatellite-stable adenocarcinomas from patients with a BMI of 24–32, and no patients relapsed after 8–19 months of follow-up. The liver metastatic sample is histologically adenocarcinoma (R1) and represents a synchronous single liver metastasis in liver segment 6 from locally advanced rectal cancer that also has one synchronous lung metastasis, staged as T3c N2 EMVI-M1. The rectal cancer patient received preoperative radiation. The metastasis was surgically resected locally from the liver before surgery was performed on the primary tumor without preoperative chemotherapy. Panel sequencing revealed no mutations in EGFR, KRAS, NRAS, BRAF, or PIK3CA (Oncomine Solid Tumor Panel Ion S5). MMR proteins are not affected, identifying the tumor as microsatellite-stable (MSI-) cancer. The samples were analyzed by a board-certified pathologist.

### Method details

#### Cell culture and treatments

The human colon cancer cell line (HCT116) and its OSE-mutant derivatives (D3 and E4) were maintained as described previously^19^. The cells were treated with either 100 μM CDK9 inhibitor (DRB; Sigma-Aldrich, D1916) for 6 hours, 2 mM Flavopiridol (Sigma-Aldrich, F3055-5MG) for 5 hours, or an equivalent amount of the solvent DMSO. Short interfering RNAs (siRNAs) were used to suppress the expression of *CCAT1* (50 nM; siCCAT1; Thermo Fisher, N510237), *CDK9* (50 nM; siCDK9; Thermo Fisher, AM51331) or *MYC* (50 nM; siMYC; Thermo Fisher, S9129) via Lipofectamine RNAiMAX transfection reagent (Life Technologies, 13778075) according to the manufacturer’s instructions. The transfected cells were maintained for 48 hours (siMYC, siCDK9) or 72 hours (siCCAT1) before harvesting.

#### Spike-in controls for RNA isolation and gating assays

To control for loss of RNA during RNA isolation, 2 ng of spike-in RNA was added to each sample before cell lysis. The spike-in RNA was generated via *in vitro* transcription from the Monster Green® Fluorescent Protein phMGFP Vector (Promega, E6421). A second spike-in was generated to control for the efficiency of the Click-iT reaction and RNA loss during nascent RNA purification. To this end, a control luciferase plasmid from the TnT Quick Coupled Transcription or Translation System (Promega, L4821) was used to generate EU-labeled spike-in RNA under the control of a T7 promoter. The *in vitro* transcription reaction was performed with a TranscriptAid T7 High Yield Transcription Kit (Thermo Scientific™, K0441) and 100 mM 5-ethynyl-uridine-5’-triphosphate sodium salt (Jena Bioscience, CLK-T08-S).

#### IncuCyte cell proliferation and confluence assay

To determine the relationship between cell proliferation and cell confluence, an equal number of WT HCT116 cells or D3 or E4 cells were seeded into a 6-well plate with the following initial cell numbers: 0.2 × 10^6^ cells/per well for achieving 10% confluence; 1 × 10^6^ cells/well for generating 50% confluence; and 2 × 10^6^ cells/per corresponding to 100% confluence. An IncuCyte S3 live-cell imaging analysis system (Sartorius) was used to determine the confluence of the cells at each time point. At the end of the experiment, RNA was extracted and quantitated for total recovery via the GFP RNA spike-in, as exemplified in **Figures 1H; S1C**.

#### Newly synthesized RNA purification

Newly synthesized RNA was labeled by incubating the cells with 0.5 mM (final concentration) 5’-ethynyl uridine (EU; Thermo Fisher, C10365) for 15 or 30 min, as indicated in the Figure legends. For analyses of transcriptional activity, the 15-min labeled cells were harvested immediately, while in the pulse chase experiment, the cells were subsequently washed with 5x PBS, followed by incubation with prewarmed normal growth medium for 1 h. Cell lysis and RNA purification were performed via an RNeasy Mini kit (Qiagen, 74104) following the manufacturer’s protocol, which included spike-in RNA. The RNA was eluted in 40 μL of nuclease-free water. EU-labeled RNAs containing EU-labeled spike-in were captured via a Click-iT Nascent RNA capture kit (Thermo Fisher, C10365) following the manufacturer’s instructions before their conversion into cDNA via a SuperScript IV VILO cDNA Synthesis Kit (Life Technology, 11766500).

#### Nuclear export assay

To determine the ratio of newly synthesized exported cytoplasmic RNAs to newly synthesized nuclear RNAs, newly synthesized RNAs were pulse-labeled with EU. After a 1-hour chase, the nuclear and cytoplasmic fractions were separated via the Ambion® PARIS™ system (Thermo Fisher, AM1921) following the manufacturer’s protocol. EU-labeled RNAs from both fractions were then purified in the presence of spike-in RNAs as above, converted to cDNA, and analyzed by qRT-PCR^18^.

#### qRT-PCR analyses

The quality of the purified RNA samples was assessed (Bioanalyzer, Agilent) before cDNA synthesis (SuperScript IV VILO cDNA Synthesis Kit, Life Technology, 11766500). All the qPCRs were performed via PowerTrack™ SYBR Green Master Mix (Thermo Fisher, A46111) on a QuantStudio™ 3 (Thermo Fisher). The linear range of amplification was confirmed by serial dilution of cDNA or sonication of genomic DNA from HCT116 cells. The primer sequences and PCR conditions are listed in the **Table S1**.

#### DNA FISH analyses

The locus-specific small DNA FISH probes were prepared from a pool of PCR products spanning 8--10 kb regions of *Hin*d III sites encompassing the *MYC* promoter and gene body (chr8:128,746,000--128,756,177) and the OSE (positioned at chr8:128,216,526--128,225,855), respectively^18^. The probes were labeled with either Cy3-dCTP (Sigma-Aldrich, GEPA53021) or Cy5-dUTP (Sigma-Aldrich, GEPA55022) via a Bioprime Array CGH Kit (Life Technologies, 18095011) and hybridized to formaldehyde-crosslinked cells as described previously^18^. To ensure that the PCR probes identified the correct genomic regions surrounding the OSE and *MYC*, we used labeled BAC clones CTD-2045B24 and CTD-3066D1 alone or in combination, as described previously^18^. Following blocking with Cot-1 DNA (Invitrogen, 15279-011), the cells were hybridized with DNA FISH probes overnight at 37°C. The cells were subsequently washed twice in 2x SSC/50% formamide at 40°C for 15 minutes and twice in 2x SSC at 40°C for 15 minutes. Finally, the cells were counterstained with DAPI solution (Thermo Fisher Scientific, 62248) and mounted in Vectashield mounting medium (Vector Labs, H-1900).

#### RNA FISH analyses

The RNA FISH analyses, which used *MYC* exon (Biosearch technologies, VSMF-2228-5) and *MYC* intron (Biosearch technologies, ISMF-2067-5) probes, were performed following the protocol from the Stellaris website. The *CCAT1* RNA FISH analyses and the *MYC* RNA FISH analyses in DRB-treated HCT116 cells were performed via PCR fragments, as described earlier^19^.

#### Immunocytochemical staining and coefficient of variation analyses

WT, D3 and E4 HCT116 cells were fixed at 50% confluence with PBS containing 1% paraformaldehyde for 10 min at room temperature, rinsed in PBS, permeabilized with 2x SCC/0.5% Triton X-100 for 10 min at room temperature, and rinsed in 2x SSC. After blocking with 10% goat serum for 1 hour at 37°C, the slides were stained with primary antibodies against c-MYC (Abcam, ab32027, 1: 50) at 4°C overnight, washed with PBS-0.05% Tween 20 four times for 15 minutes each, and then blocked with goat serum before incubation with secondary antibodies (Alexa Fluor 594-conjugated anti-rabbit or Alexa Fluor 546-conjugated anti-rabbit) at 37°C for 2 hours. After being washed with 1x PBS containing 0.05% Tween 20, the cells were stained with DAPI for 5∼10 min. Immuno-stained cells were mounted in Vectashield mounting medium (Vector Labs, H-1900), covered with coverslips and visualized with a wide-field microscope (Leica DMi8). Nuclei were identified *via* the DAPI channel, and MYC intensity was quantified specifically within nuclear boundaries, excluding the cytoplasmic signal, via CellProfiler 4.2.8 using optimized parameters identified in **Figure S2H**.

Thin sections (10 μm) of CRC samples were stained with anti-MYC primary antibody (Abcam, ab32027, 1:50)and PAN cytokeratin antibody (Invitrogen, 53-9003-8, 1:100 dilution) overnight at 4°C in an antibody mixture (Life Technologies, #003218) after initial blocking in 10% goat serum for 1 h at 37°C. The slides were washed with PBS-0.05% Tween 20 four times for 15 minutes each, followed by blocking with 10% goat serum for 1 h at 37°C before incubation with secondary antibodies conjugated to Alexa Fluor 594 or 488 and DAPI counterstaining. Imaging was performed on a Leica fluorescence microscope under identical conditions for all the samples. The combined immunostaining and DNA FISH analyses followed the protocols detailed above. To compare the same motifs, images of selected areas were captured both before and after the DNA FISH analyses. Cells that were not represented by complete nuclei upon 3D reconstruction were excluded from the analyses.

#### Chromatin-immunoprecipitation (ChIP)

The samples were fixed with 1% PFA for 10 min at RT. Formaldehyde was quenched with 0.125 M glycine for 10%; the cells were then collected by scraping into new 0.125 M glycine containing protease inhibitor cocktail (PIC, Roche Merck 4693132001), and the pellet was collected by centrifugation at 2000 × g for 4 min at 4°C. After approximately 10^7^ cells were lysed (50 mM Tris-HCl pH 8, 10 mM EDTA, 1% SDS, 1X PIC) 10’ on ice, the chromatin was sonicated with Covaris for 10 min at 4°C to produce fragments between 600 bp and 1500 bp (20μl of sonicated chromatin was treated with RNase and proteinase K O/N at 65°C, purified in phenol:chloroform, quantified by Nanodrop and run on a 1% agarose gel to check sonication efficiency). The sonicated chromatin was then centrifuged at 10’ and 4°C at 1000 rpm to remove insoluble material, and the supernatant was transferred to a fresh tube. A total of 30μg of diluted chromatin in IP buffer (15 mM Tris pH 7.5, 1 mM EDTA, 150 mM NaCl, 0.5% Triton X-100, 1X PIC) was incubated with 4μg of anti-RNAPII-CTD-phospho-S5 antibody (Abcam ab5408) or anti-unmodified-RNAPII-CTD (Santa Cruz sc-56767) O/N at 4°C with rotation. The antibody-chromatin complex was then incubated with 15μl of blocked (with 1 mg/ml BSA and 0.1μg of yeast tRNA in IP buffer for 30’ at RT) Pierce protein A/G magnetic beads (Thermo Scientific 88802) at 4°C for 2 h with rotation. The bead-antibody-chromatin complex was washed twice at 10°C at 4°C with 1 ml of each of the 4 wash buffers (WBs) (WB 1: 20 mM Tris pH 7.5, 2 mM EDTA, 150 mM NaCl, 0.5% Triton X-100, 0.1% SDS). WB 2: same as WB1 but with 500 mM NaCl. WB3: 10 mM Tris pH 7.5, 1 mM EDTA, 250 mM LiCl, 1% Triton X-100, 1% sodium deoxycholate, and 0.1% SDS. WB4: 10 mM Tris pH 7.5, 1 mM EDTA). Finally, the immunoprecipitated DNA and 10% input material were reverse-crosslinked (0.2 M NaCl, 50 mM Tris pH 7.5, 1 mM EDTA, 1% SDS, 200μg/ml proteinase K, Sigma P2308) O/N at 65°C, purified by ChIP clean and concentrator columns (Zymo Research D5205) according to the manufacturer’s protocol and eluted in 30μl. qPCR was performed to compare the enrichment signal of the ChIP to that of the input.

#### ChrISP

BAC DNA FISH probes targeting the OSE (BAC clone CTD-2045B24) and *MYC* (BAC clone CTD-3066D1) regions were labeled with digoxigenin-11-dUTP (Roche, 11573152910) and biotin-16-dUTP (Roche, 11093070910), respectively, via the BioPrime® Array CGH labeling kit (Thermo Fisher Scientific, 18095-011). Human breast cancer tissue cryosections were cut at 8 µm thickness and fixed with 2% paraformaldehyde (PFA) in PBS, followed by permeabilization in 2×SSC containing 0.5% Triton X-100 for 10 min at RT. After being rinsed with 2× SSC, the tissue sections were denatured in 2×SSC with 50 % formamide at 80°C for 40 min and subsequently incubated in ice-cold 2×SSC for 5 min. After being blocked with Cot-1 DNA at 37°C for 30 min, the FISH probes were added together with human Cot-1 DNA (Invitrogen, 15279–011) and hybridized to the slides in a buffer containing 2× SSC,50% formamide and 10% dextran sulfate overnight at 37°C. The slides were then washed twice with 2×SSC in 50% formamide for 15 minutes at 40°C, followed by washing with 2×SSC for 15 minutes at 42°C twice. For ChrISP analysis between the DIG-labeled OSE probes and the nuclear pore component NUP133, a Tyramide signal amplification step (Biotin XX Tyramide SuperBoost™ Kit, goat anti-rabbit IgG, LifeTechnologies, B40921) was included to increase the concentration of biotin molecules in the vicinity of the NUP133 epitopes. The TSA reaction was performed as described in the manufacturer’s protocol.

In brief, after DNA FISH (as above) and immunostaining with a primary antibody against NUP133 (Abcam, ab155990; diluted 1:200) overnight at 4°C, the tissue sections were washed with 0.05% Tween 20 in PBS and blocked for 1 hour at room temperature with 10% goat serum, followed by incubation with a poly-HRP-conjugated secondary antibody for 1 hour at room temperature. After a further washing step in PBS, the cells were treated with Tyramide working solution for 5 min, washed again with PBS and incubated with an antibiotin antibody (Cell Signaling, 5597S; diluted 1:200) and an anti-digoxigenin antibody (Roche, 11333062910; diluted 1:200) overnight at 4°C. After an overnight washing step in 0.1×SSC at 37°C, the ChrISP assay was performed as previously described^18^. Negative controls omitting the mouse secondary ChrISP antibody (“R-” control) were included in parallel for each experiment. Finally, the tissue sections were counterstained with DAPI (Thermo Fisher Scientific, 62248) and mounted with Vectashield mounting medium (Vector Laboratories, H-1900).

#### Wide-field microscopy

Cell imaging and 3D optical sectioning were performed via a Leica DMi8 microscope fitted with an HC PL APO 63X NA 1.4 oil objective and a DFC9000 camera. A Thunder Imaging System (Leica Microsystems) was utilized, employing the Instant Computational Clearing Method to enhance image clarity. Z-stack acquisition was carried out with software-optimized interval settings along the z-axis. Image analysis was conducted via Imaris or Leica Application Suite X (LasX) software. Given the resolution constraints of the fluorophores, with CY3 resolving at 239.6 nanometers, the distance data were stratified using 240 nanometers as the initial threshold.

### Computer simulation, quantification and statistical analyses

Statistical analyses were performed via GraphPad Prism and Microsoft Excel. The mean nuclear MYC intensity per cell in tumor specimens was calculated and plotted via Prism, and heterogeneity across nuclei was quantified by determining the coefficient of variation (%CV) within each replicate. Comparisons of mean intensities across multiple groups were assessed via one-way ANOVA, followed by Tukey’s post hoc test when applicable, with exact p values displayed in the corresponding bar graphs.

For pairwise comparisons between genotypes (WT *versus* mutant), two-tailed Student’s t tests were applied—unpaired t tests for independent groups and one-sample t tests where appropriate to assess deviation from a normalized reference. All t tests assumed a normal distribution, equal variances, and independence of observations and were two-sided unless otherwise stated. Pearson correlation coefficients were calculated for association analyses. For categorical comparisons, such as spatial RNA/DNA FISH patterns, significance was evaluated via Fisher’s exact test. The results are presented as the means ± standard deviations unless otherwise specified.

To model the contribution of MYC gating to total *MYC* mRNA levels, we incorporated the values of the transcription level, *MYC* mRNA nuclear export rates and nuclear and cytoplasmic *MYC* mRNA degradation rates into a previously published algorithm^18^.

The microscopic data were curated in Excel and processed in R 4.3 via the tidyverse, readxl, car, ineq, entropy, philentropy, mclust, diptest, transport, ggpubr, ggplot2, and openxlsx packages. A unified R workflow (run_heterogeneity_analysis) was written and applied to quantify expression heterogeneity. The metrics included the coefficient of variation, Gini coefficient, and distribution-based divergence measure (Jensen–Shannon divergence). Fligner–Killeen and Levene’s tests were used to assess the equality of variances, and Hartigan’s dip test combined with Mclust mixture modeling was used to evaluate distributional multimodality. For tumor analyses, per-cell data were first annotated into manually defined topological classes. All tests were two-sided, and *p* < 0.05 was considered statistically significant. Heterogeneity in nuclear MYC intensity and related parameters was quantified via multiple complementary indices capturing different aspects of population variability:

Coefficient of variation (CV): measured normalized dispersion (σ/μ), indicating the relative spread of MYC intensity values across cells within a group. A higher CV reflects broader variability in MYC expression between individual nuclei.

Gini coefficient: This coefficient measures inequality in intensity values, analogous to income inequality. Gini = 0 indicates uniform expression; Gini → 1 indicates extreme disparity dominated by a small subset of high-MYC cells.

Fligner–Killeen and Levene’s tests: evaluate the equality of variances across groups or clusters. The significant results indicate that one or more groups exhibit distinct expression variability. Hartigan’s dip test and multiplex mixture modeling: identify multimodal distributions, revealing subpopulations with distinct MYC expression states within a group or cluster.

All reagents are listed in **Key Resource Table** with included RRID numbers when available.

## Resource availability

### Lead contact

Requests for information should be directed to and will be fulfilled by the lead contact, Anita Göndör

### Materials availability

All unique/stable reagents and data generated in this study are available from the lead contact The data code used to analyze heterogeneity metrics is available upon request

## Acknowledgements

We gratefully acknowledge the permission from the patients to use their tumor material for analyses. This work was supported by the Swedish Research Council (VR; 2021-02570; 2019-01553; 2024-03026), the Swedish Childhood Cancer Fund (PR2020-0162; PR2023-0127), the Swedish Cancer Society (19 0381 Pj; 22 2340 Pj; 22 2175), the Lundberg Foundation (2018-0138), The KA Wallenberg Foundation (KAW 2017.0077), Karolinska Institutet, the Novo Nordisk Foundation (NNF16OC0021512), The Cancer Society in Stockholm (Cancerföreningen in Stockholm, 2018; 2019), the Norwegian Cancer Society (247350), Helse Sør-Øst RHF (2018054, 2025002) and AHUS strategic grants.

## Author contributions

CG, MM, AC, FBC, BS, IT, YL, YvE, WF, TM, JPL, RK, PC - RNA/DNA FISH analyses; CG, AC, MM, IC - export assays; CG - DRB experiments; MM, IT, YvE - Flavopiridol/siCDK experiments; MM, ChIP experiments; JPL, BS - ChrISP experiments; MM, WF, CG - siRNA experiments; PM – RNA decay analyses; DT, YL, YvE, TM – MYC protein expression; SM, AHR, JH, NG and MG – collection and characterization of the tumor tissue samples; NK, MT, AG, RO – data and statistical analyses; AG, RO, CG, MM wrote the manuscript; AG, RO, AHR, JH, MG – secured funding; AG – overall experimental design and planning.

## Supplementary Figures and legends

**Figure S1.**
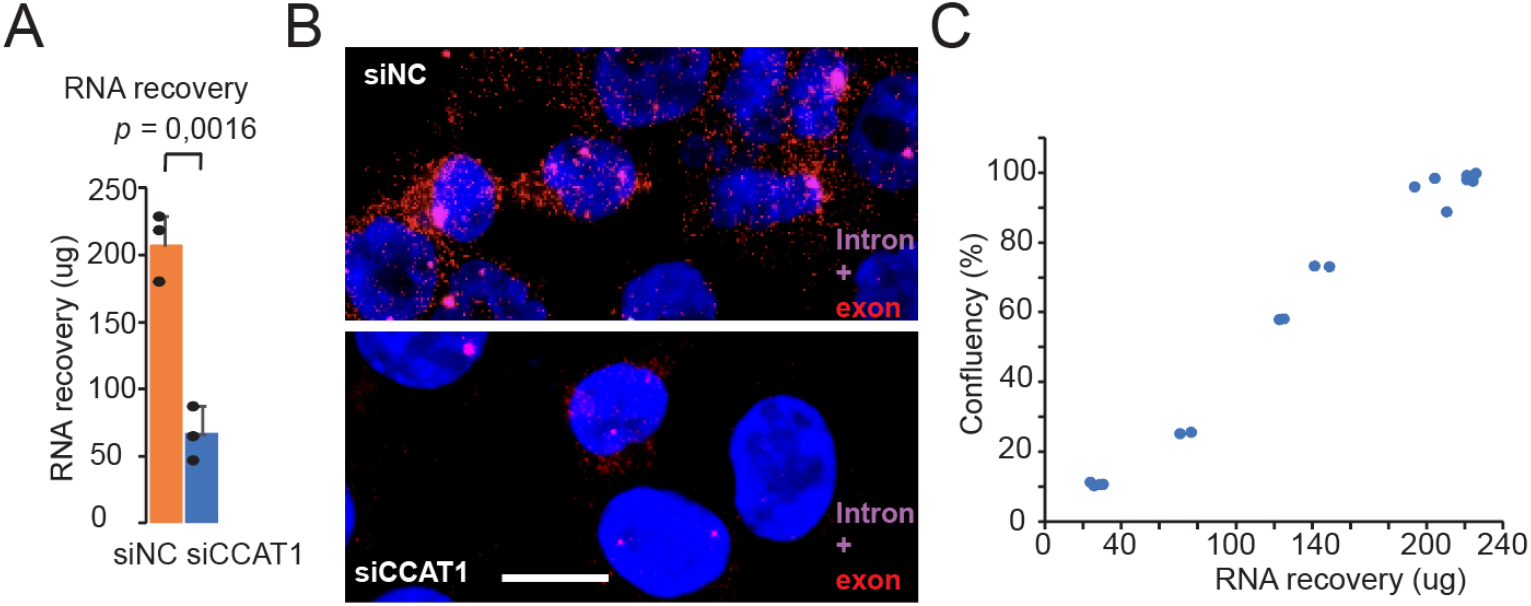
MYC gating is regulated by CCAT1 and cell confluence. A) Total recovery of RNA, normalized to the GFP spike-in RNA, in the control (siNC) and siCCAT1-treated samples. B) Representative images of RNA FISH analyses of siNC- or siCCAT1-treated cells using Stellaris *MYC* exon and intron probes. C) Determination of cell confluence via the recovery of extracted RNA. The cells were seeded at different densities and collected at indicated cell confluences, as determined by the IncuCyte. The cell extract was spiked with GFP RNA to compensate for any RNA loss during extraction. The *P* value was determined by the two-tailed, unpaired Student’s t test. Bar = 10 μm.

**Figure S2.**
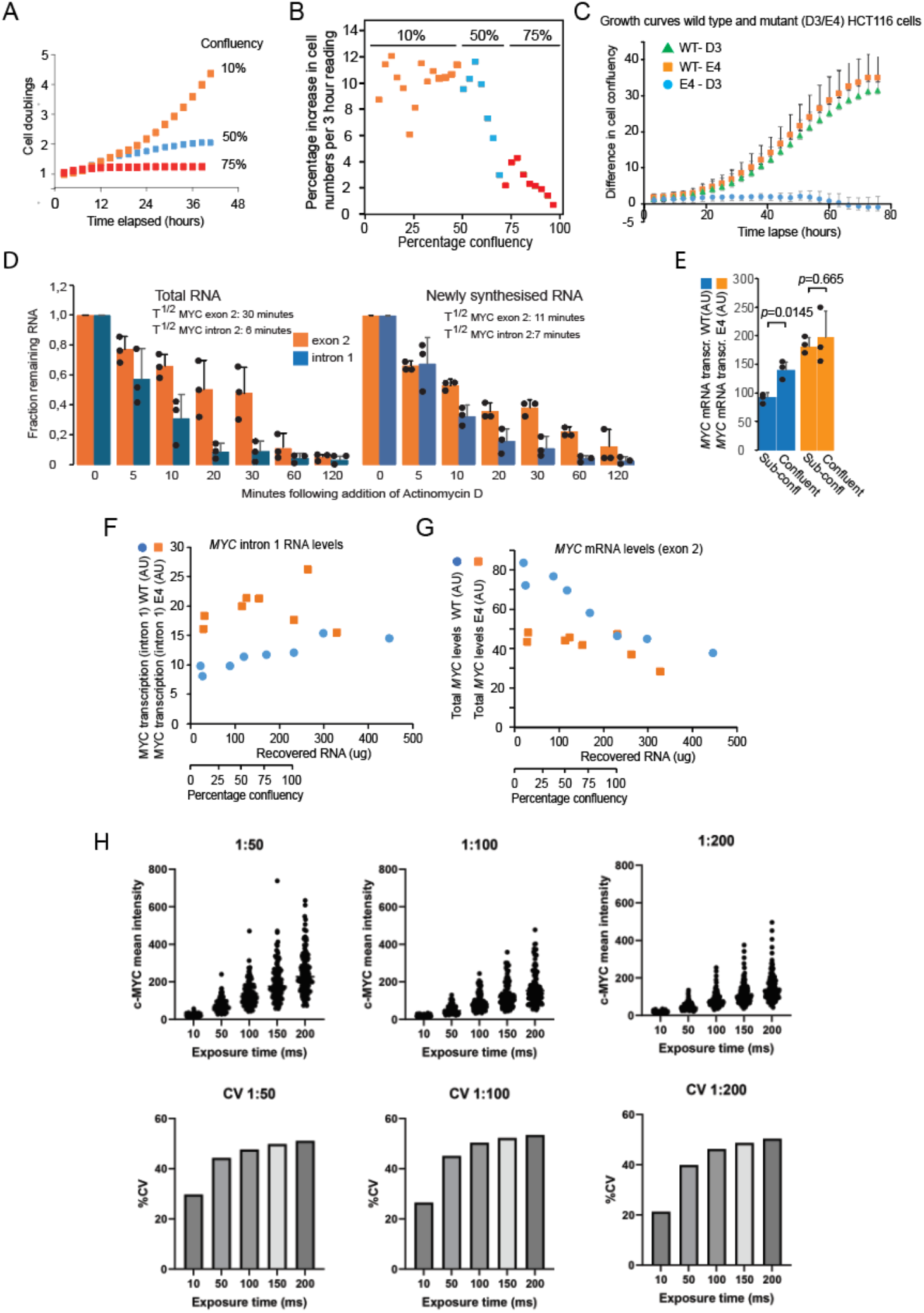
MYC gating drives both exponential cell proliferation and MYC expression heterogeneity. A) Cell growth curves of cells seeded at the confluence indicated in the image. B) Data from (A) were stratified to show cell proliferation at 3-hour intervals. The first three data points of each set of data were removed from the image due to trypsinization effects (see text for more information). C) WT HCT116, E4 and D3 (independent OSE-mutant HCT116 clones) cells were seeded at approximately 10% confluence and then incubated for 72 hours in the IncuCyte system. The differences in cell proliferation are depicted on the y-axis. The data shown are the means of two independent experiments. D) *MYC* mRNA decay kinetics in total RNA (left) and newly synthesized RNA fractions (right) following the addition of actinomycin D. The data from three independent experiments were normalized to the 0-hour time point. E) MYC transcription rates determined by intron 1 signals, normalized to spike RNAs for the calculation of total RNA recovery and to PCR efficiencies of cells harvested at <40% and 100% confluence, in three independent experiments. *MYC* transcription levels (F) and *MYC* mRNA accumulation (G), normalized to RNA input, in WT HCT116 cells and E4 cells carrying a mutated CTCF binding site within the intron of *CCAT1* are depicted in relation to cell confluence. H) Simulation of the contribution of gating to total *MYC* mRNA accumulation (right, diagram) by integrating mRNA decay data, nuclear export rates and transcription rates, as previously described^18,19^. The blue and red lines depict the values obtained via nuclear export rates of 8 (gated) and 1 (nongated), respectively. P1 = 8.0897E-05 (two-tailed, unpaired Student’s T-test). The decay data (D) and transcriptional and nuclear export rates were used to simulate the contribution of *MYC* gating to overall levels of *MYC* mRNA accumulation at low cell confluence. H) Optimization of the MYC staining protocol for obtaining reproducible coefficients of variation.

**Figure S3.**
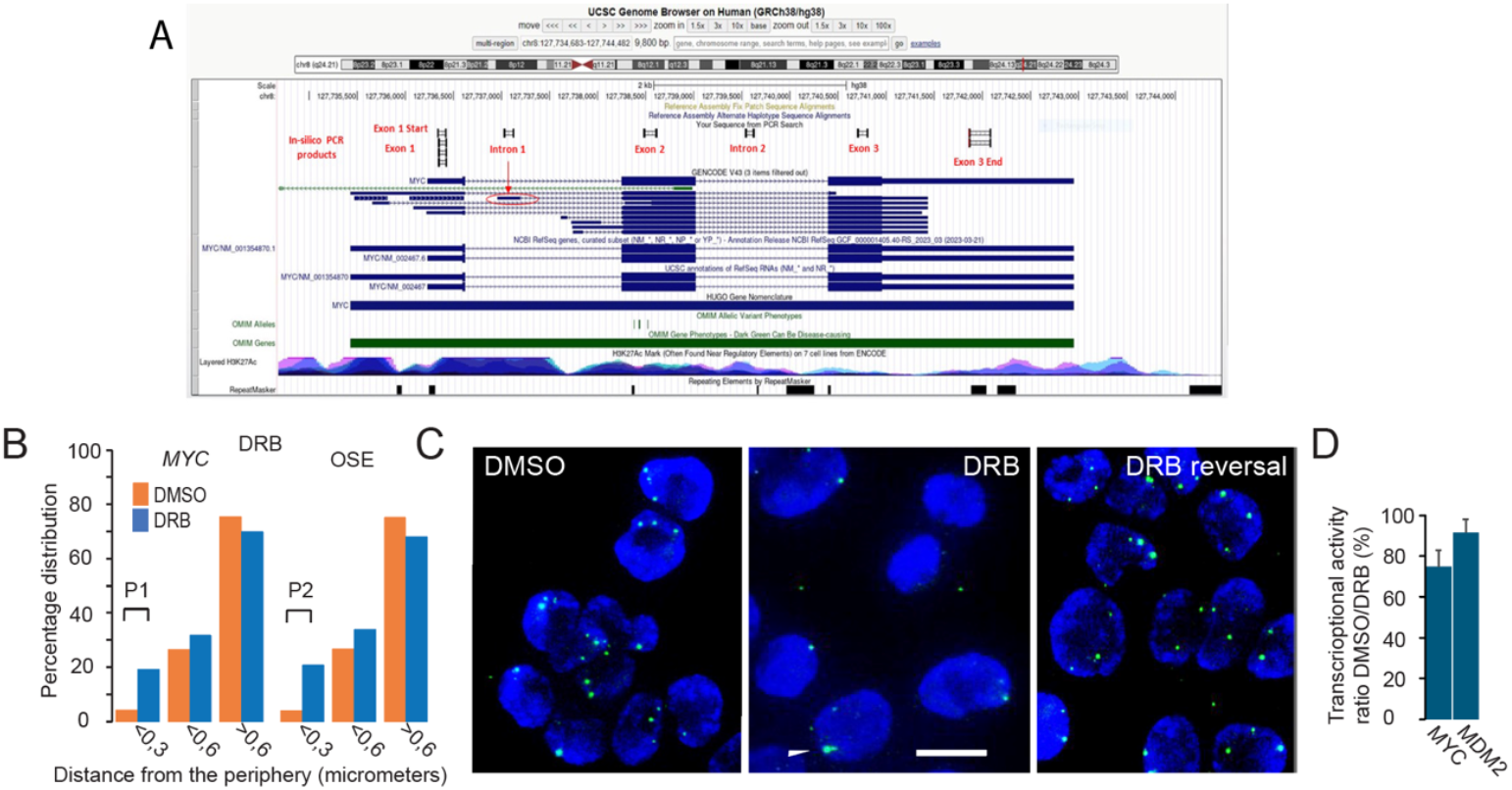
Inhibition of transcriptional elongation promotes MYC migration to the nuclear periphery. A) Map of PCR primers. B) DRB treatment increased the presence of *MYC* and OSE alleles at the nuclear periphery in WT HCT116 cells. The results are the average of two independent experiments (fixed at 50% confluence), with a total of 968 alleles counted. P1 = <0.0001; P2 = 0.0013. C) RNA FISH analyses of *MYC* expression via PCR probes covering the entire *MYC* gene. D) Transcriptional rates of *MYC* and *MDM2* upon DRB reversal at 0 minutes. The results are the average of three independent experiments unless otherwise indicated. The *P* values were determined by Fisher’s exact test (A). Bar = 10 μm.

**Figure S6.**
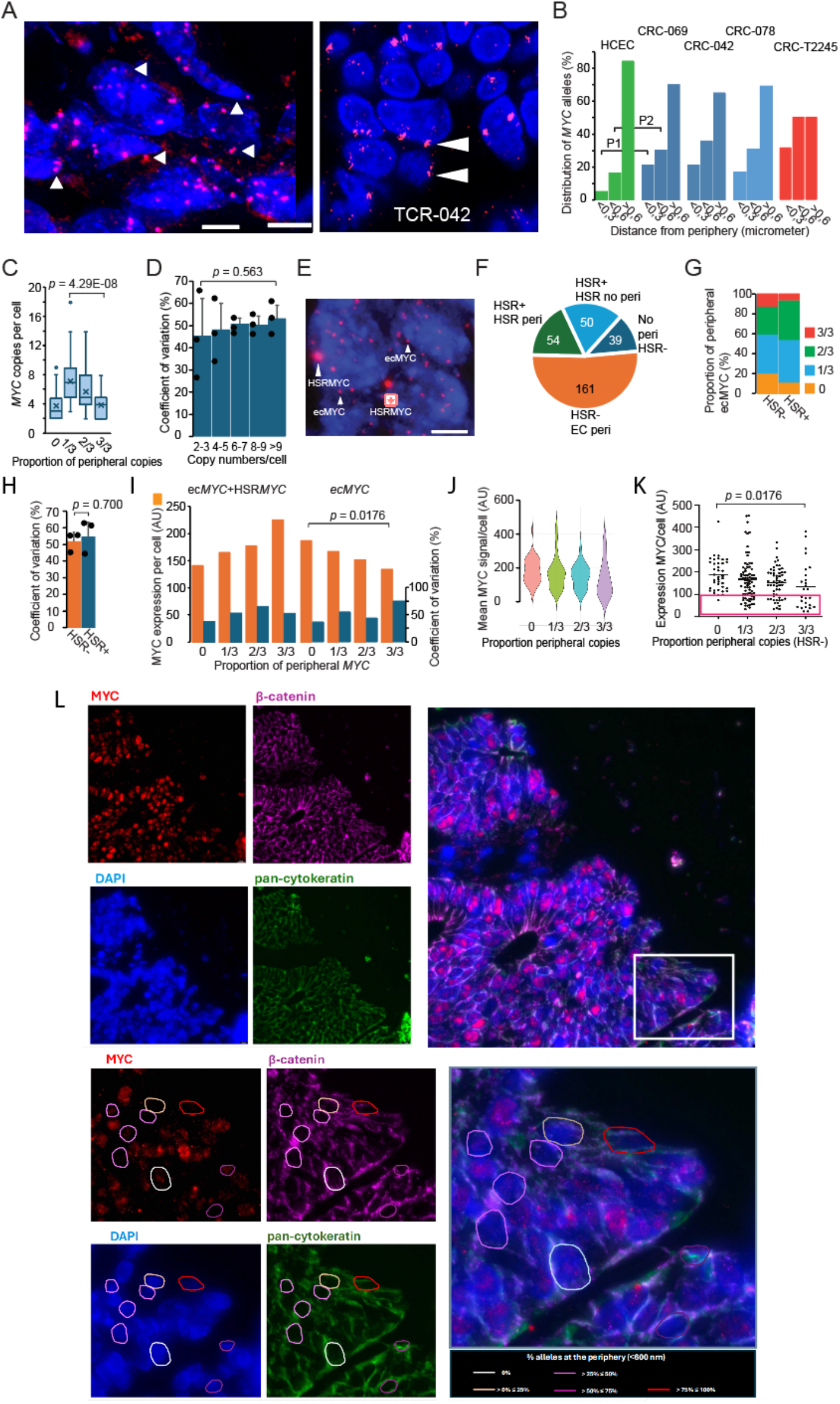
Hallmarks of MYC gating correlate with MYC expression heterogeneity in tumors. A) Representative images of the distribution of *MYC*-containing HSRs and ecDNA in metastatic (liver, left) and primary (right, confocal plane) colorectal cancers. Arrows denote some of the multiple peripheral HSRs. F) Quantification of the spatial distribution of *MYC* alleles in three primary CRC samples and one liver metastasis of CRC. The data, which are based on a total of 504 (CRC-069), 758 (CRC-042) and 687 (CRC-078) alleles, are compared to 324 alleles from primary cultures of human colon epithelial cells (HCECs)^18^. P1 = <0.0001, P2 = < 0.0001. C) Box plot analyses of the number of *MYC* copies per cell stratified according to the frequency of their peripheral distribution. D) Coefficient of variation of the mean intensity of nuclear MYC staining determined in cells with 2-3, 4-5, 6-7, 8-9, or >9 copies of *MYC*. The data are based on a total of 304 cells (1242 *MYC* alleles). E) Representative image from tumor CRC-042 showing cells displaying extrachromosomal *MYC* copies (ec*MYC*) with or without *MYC* amplified in homogenously staining regions (HSR*MYC*). F) Pie diagram visualizing the number of tumor cells in each category (HSR-peri = cells HSR*MYC-/*peripheral ec*MYC*; HSR- = cells HSR *MYC-/*no peripheral ec*MYC*; HSR+ no HSR peri = HSR+/no peripheral HSR*MYC*; HSR+ HSR peri = HSR+/peripheral HSR*MYC*). More than 95% of all signals included both *MYC* and distal OSE, thus encompassing > 500 kb domains. G) Bar diagram showing the distribution of cell populations based on the proportion of peripheral ec*MYC* copies in only HSR- and HSR+ cells. H) Analyses of MYC expression heterogeneity (CV) in HSR- or HSR+ cells from three different CRC samples. I) Comparison of the expression levels and CV values of the mean nuclear MYC levels between the HSR- (201 cells) and HSR+ (103 cells) populations. The data were stratified according to the proportion of peripheral *MYC* copies. Violin (J) and scatter (K) plots of MYC expression profiles in HSR-negative cells stratified by the proportion of peripheral ec*MYC* copies. The red box in panel H represents an increasing proportion of cells with low MYC expression (<100 AU) correlating with an increasing proportion of peripheral ec*MYC* copies. L) Cytokeratin-stained CRC tumor cells in thin sections. The boxed area represents one of many motifs that were examined and that contains a spectrum of MYC expression levels and how they correlated with peripheral ecMYC copies (color-coded to indicate the degree of peripheral distribution). The bar corresponds to 10 μm. The *P* values were determined by Fisher’s exact test (B) or the unpaired, two-tailed Student’s t test (C, D, H, I, K).

